# Genetic program activity delineates risk, relapse, and therapy responsiveness in Multiple Myeloma

**DOI:** 10.1101/2020.04.01.012351

**Authors:** Matthew A. Wall, Serdar Turkarslan, Wei-Ju Wu, Samuel A. Danziger, David J. Reiss, Mike J. Mason, Andrew P. Dervan, Matthew W.B. Trotter, Douglas Bassett, Robert M. Hershberg, Adrián López García de Lomana, Alexander V. Ratushny, Nitin S. Baliga

## Abstract

Despite recent advancements in the treatment of multiple myeloma (MM), nearly all patients ultimately relapse and many become refractory to their previous therapies. Although many therapies exist with diverse mechanisms of action, it is not yet clear how the differences in MM biology across patients impacts the likelihood of success for existing therapies and those in the pipeline. Therefore, we not only need the ability to predict which patients are at high risk for disease progression, but also a means to understand the mechanisms underlying their risk. We hypothesized that knowledge of the biological networks that give rise to MM, specifically the transcriptional regulatory network (TRN) and the mechanisms by which mutations impact gene regulation, would enable improved predictions of disease progression and actionable insights for treatment. Here we present a method to infer TRNs from multi-omics data and apply it to the generation of a MM TRN that links chromosomal abnormalities and somatic mutations to downstream effects on gene expression via perturbation of transcriptional regulators. We find that 141 genetic programs underlie the disease and that the activity profile of these programs fall into one of 25 distinct transcriptional states. These transcriptional signatures prove to be more predictive of outcomes than do mutations and reveal plausible mechanisms for relapse, including the establishment of an immuno-suppressive microenvironment. Moreover, we observe subtype-specific vulnerabilities to interventions with existing drugs and motivate the development of new targeted therapies that appear especially promising for relapsed refractory MM.

## Introduction

Multiple Myeloma (MM) is a cancer of malignant plasma cells in the bone marrow (BM) that has a prevalence of approximately 86,000 new cases per year.^1^ Several clinical subtypes of MM have been established on the basis of characteristic cytogenetic features, including various translocations, gain or loss of chromosomal arms, deletion of specific chromosomes, and hyperdiploidy.^2–4^ Accordingly, MM is a complex disease of great heterogeneity that exhibits subtype-specific drivers of progression.^5,6^ Efforts to better characterize the biology and therapeutic vulnerabilities of MM have increased exponentially in recent years, as can be seen from the number of research articles, clinical trials, and public availability of matched genomic, transcriptomic, and patient data. However, the myriad combinations of chromosomal aberrations and somatic mutations, coupled with the complex dependence of MM progression on the BM microenvironment, have precluded a mechanistic understanding of the disease on a patient-specific level.

The 5-year survival rate of MM is approximately 50%^7^, which is nearly double what it was in the late 1980s.^8^ The reason for the improved outcomes is a series of therapeutic advances that include the introduction of autologous stem cell transplants (ASCT), immunomodulatory drugs (IMiDs), and proteasome inhibitors (PIs), all of which are now first-line therapies.^7–9^ Despite the success of these treatments in extending the overall survival (OS) by 6-10 years, depending upon age at diagnosis, most patients eventually relapse and become refractory to the therapies that they had received.^10–12^ The mechanisms by which patients develop resistance to these therapies are only partially understood and are thought to depend critically upon the BM microenvironment.^3,13–15^ Nonetheless, many new therapeutics with diverse mechanisms of action are under development and showing promise in clinical trials against relapse refractory multiple myeloma (RRMM).^8^ These include next-generation IMiDs and PIs, as well as histone deacetylase (HDAC) inhibitors, monoclonal antibodies (mAbs), and immunotherapies. Currently, the optimal combination and sequence of these therapies is unknown, both at baseline and relapse.^16^

If the disease biology of individual patients can be sufficiently well characterized from experimental assays like RNA-sequencing and cytogenetics, it is conceivable that they can be assigned the best available therapies and manage their cancer like a chronic illness with much-improved outcomes. However, a detailed map of the underlying biology of MM is necessary to translate the data collected from a patient into personalized recommendations for therapy. The development of such a map is complicated by the great degree of heterogeneity MM exhibits, including subtypes at the levels of gene expression, gene mutations, chromosomal abnormalities, and clinical outcomes. Before we can establish an era of personalized medicine for all MM patients, we must understand how the subtypes at these different levels relate to one another mechanistically, and which of these features are most important for determining the risk of disease progression. Moreover, we must characterize the biological changes that drive escape from therapy and the onset of relapse refractory disease. Once we understand the subtype-specific drivers of disease progression and biology of relapse, we can rationalize and test which therapies are most appropriate for which patient subtypes.

We hypothesized that a causal mechanistic (CM) transcriptional regulatory network (TRN) would provide a robust framework to establish a biological context of MM subtypes and reveal actionable insights into patient-specific susceptibilities to respond or escape from different therapies. The advantage of a CM TRN is that it can reveal putative mechanisms of gene regulation and how different mutations and chromosomal aberrations dysregulate these processes leading to hallmarks of cancer.^17^ Patterns in the CM TRN can then be related to clinical outcomes in order to elucidate the biological context of a heterogeneous disease. Thus, the different levels of MM heterogeneity can all be linked to one another in a CM TRN. However, there are several challenges to the development and application of CM TRNs, including spurious correlations that arise from the high dimensionality of gene expression data, difficulty of detecting rare features such as condition-specific regulatory mechanisms, complexity of inferring causal events, and the requirement of efficient computational algorithms.^18–21^ Moreover, there is no generally accepted protocol to infer which features of the network are activated or deactivated in an individual patient. This is of critical importance, because the CM TRN represents possible biological mechanisms across all subtypes of a disease, such that each individual will only exhibit activity in a subset of the network features.

Although gene expression networks have previously been derived to study MM^22–24^, a CM TRN that elucidates causal flows from mutations to regulators to co-regulated genes across MM subtypes has not yet been established. In this work, we present a novel method called Mechanistic Inference of Node-Edge Relationships (MINER) to construct a CM TRN from multi-omics and clinical outcomes data, infer patient-specific network activity, and identify subtype-specific mechanisms that are likely to predispose resistance or susceptibility to a given therapy. We apply this method to multi-omics data from multiple myeloma patients in order to better characterize the disease, with the specific goal of elucidating the underlying biology of high-risk clinical subtypes and the changes that occur at relapse.

## Results

### MINER pipeline infers causal-mechanistic transcriptional regulatory network (CM TRN) of MM

We developed the MINER pipeline to infer transcriptional regulatory networks from gene expression data and apply them to the characterization and prediction of phenotypes. MINER builds upon our previous work with the SYstems Genetics Network AnaLysis (SYGNAL) pipeline insofar as it enables the same core functionalities of mechanistic and causal inference, but does so with a new suite of algorithms that enable new applications in the network-based prediction of clinical outcomes (**Fig. 1**)^17^. Inference of the TRN begins by clustering gene expression data into coherent sets of genes that share a binding site for a transcription factor or miRNA according to gold-standard binding-site database information (*e*.*g*., *Transcription Factor Binding Site Database - TFBSDB*). By default, the cluster of genes and the corresponding regulator(s) must also be correlated (or anticorrelated) to one another, however this restriction can be lifted if it proves too stringent, for example in single cell analysis. The combination of a coherently expressed set of genes and the associated regulator(s) whose binding site(s) they share represent discrete units, called *regulons*, from which the TRN is assembled. Once the regulons have been discovered, a novel causal inference algorithm (see Methods) identifies statistically significant links between putative causal events (somatic mutations, chromosomal translocations, etc.) and the activity levels of regulators and co-regulated gene sets. Once a MINER TRN has been inferred from the data of a patient cohort, new samples can be analyzed to uncover the disease-relevant modules that are over- or under-active in an individual patient.

**Figure 1.**
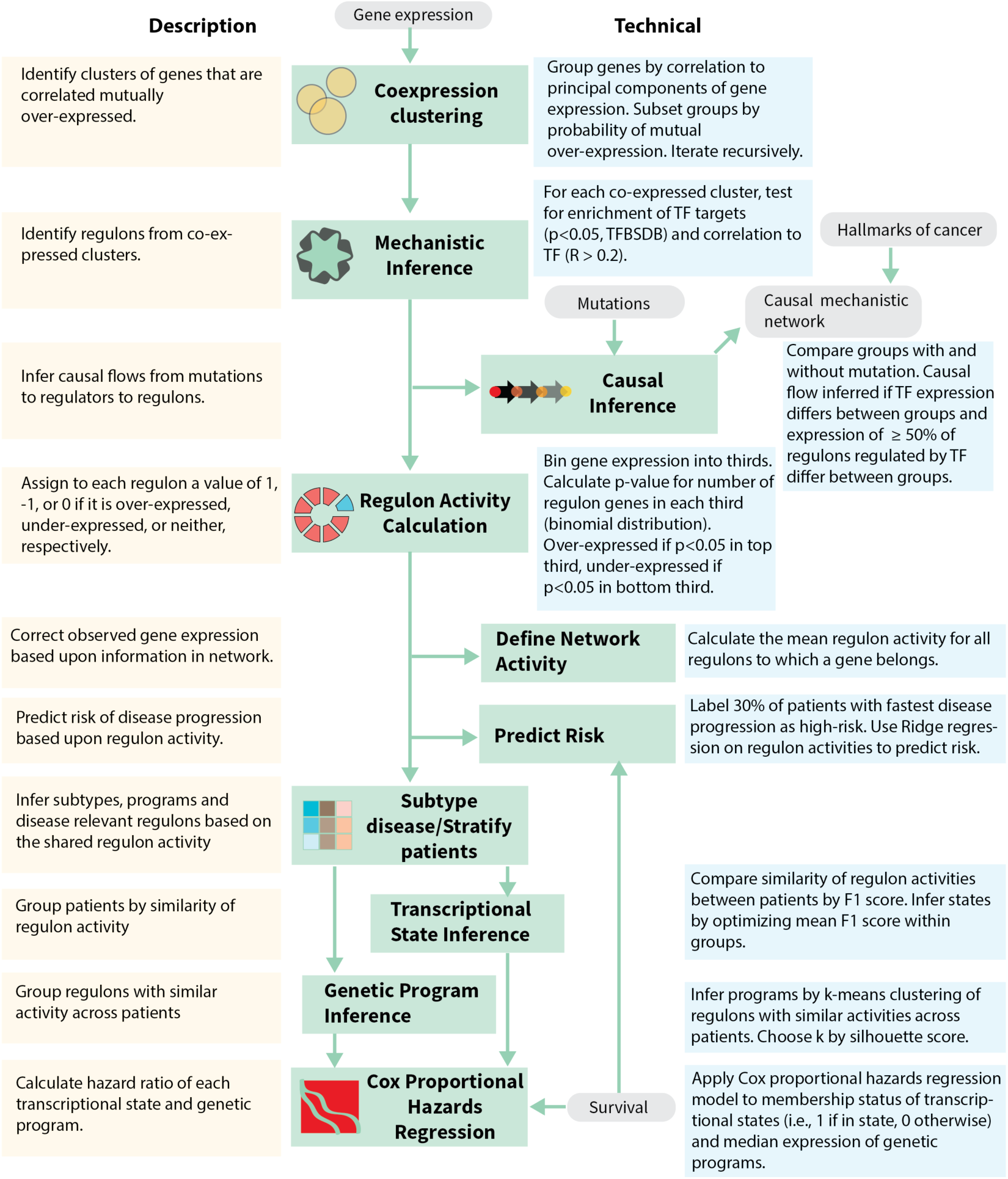
Mechanistic Inference of Node-Edge Relationships (MINER). MINER applies a gene expression clustering algorithm and gold-standard gene interaction databases to infer sets of co-regulated genes called regulons. Each regulon has an associated regulator and edge direction (i.e., activation or repression), as well as an activity in each sample (e.g., over-expressed, under-expressed, etc.). When the coordinated activity of regulons changes in the context of a mutation, a causal relationship is inferred through the associated regulator. A causal and mechanistic transcriptional regulatory network is generated by evaluating the influence of all potential causes (e.g., mutations, translocations, copy-number abnormalities, etc.) on all regulons. Prediction of phenotypes such as responsiveness to therapy or risk of disease progression is achieved by training machine-learning algorithms on regulon activities. The predictive signatures are then placed into a meaningful biological context by identifying their associated mechanisms and putative causes within the network.

We identified the Multiple Myeloma Research Foundation (MMRF) Interim Analysis 12 (IA12) dataset as the best publicly available dataset for construction of the MINER TRN (see Methods). Somatic mutation and cytogenetic calls were used as provided in the clinical data table files, and the RNA-seq translocation calls were used as provided. However, quality control of the gene expression data revealed that extensive normalization was necessary, as the most highly expressed transcripts cause an effective under-sampling of all other transcripts (**Fig. S1**). We accounted for this by applying a modified version of trimmed mean of M values (TMM) normalization^25^ that also employs quantile normalization to mitigate batch effects from composition bias (see Methods).

Application of MINER to the pre-processed MMRF IA12 data successfully generated a CM TRN of MM. The network features 15,192 genes partitioned into 1,233 co-expression clusters (i.e., without inferred co-regulation), 8,549 genes partitioned into 3,203 co-regulated modules (called *regulons* herein) that are regulated by 392 unique transcription factors, and 124 causal drivers, including somatic mutations, translocations, and cytogenetic abnormalities (complete network provided in Supplementary File S1). In total, the MINER network comprises 13,587 unique CM flows – links from mutations to regulators to co-regulated genes. We note that miRNA-seq was not performed in the MMRF CoMMpass study, so the inference of miRNA regulation was limited to analysis of target gene expression (see Methods) and was therefore less robust than transcription factor analysis. The results of our miRNA inference are provided in the Supporting Information, but the remainder of this manuscript will focus on regulation mediated by transcription factors. Every mutation or chromosomal aberration that occurred in at least 2% of the patient population is represented within the network, such that its inferred causal effect on the regulation of gene expression is described. This includes recapitulation of the known transcriptional effects, such as the up-regulation of NSD2 by t(4;14), MAF by t(14;16), and CCND1 by t(11;14). In addition to the raw network files, we have developed an interactive web portal to facilitate investigation of the CM TRN.

### CM TRN recapitulates biology of MM and proposes downstream effects of driver mutations

We investigated the literature for evidence of the inferred causal and mechanistic relationships presented herein. First, we subset the network to focus on regulons that were at least moderately associated with risk of disease progression (i.e., |Cox hazard ratio (HR)| > 2), then searched for PubMed abstracts reporting these driver mutations, regulators, or genes as implicated in multiple myeloma or other cancers. We confirmed that at least 107 of the 109 mutations in this network had been previously associated with cancer, and 34 had been specifically associated with MM. As we incrementally increased the minimum |HR| threshold from 2 to 6, the network became restricted to higher-risk regulons and the percent of our network inferences that were validated as associated to MM in the literature monotonically increased (|HR| > 2: 31%; |HR| > 3: 33%; |HR| > 4: 36%; |HR| > 5: 41%; |HR| > 6: 46%). This is consistent with an expected bias that literature abstracts preferentially report findings that associate to risk when discussing specific mutations or genes in the context of MM. In other words, when we consider the highest risk elements of our network, we find that the driver mutations inferred by our unsupervised method are strongly corroborated as associated with MM. This not only suggests that the network recapitulates the known biology of MM, but that there is a great deal of relevant information in the network that does not exist in the literature. For example, the association of the mutations in our network with other cancers in the literature implies a larger set of oncogenic driver mutations that were not previously described for MM. Most importantly, this CM TRN provides mechanistic links between the many important but disconnected features in the literature by means of causal flows from mutations, through regulators, to co-regulated gene sets. Because of this, we can associate biology, rather than correlation alone, to risk of disease progression.

### Network quantization enables patient-specific network analysis

We developed a new approach to quantifying the activity of the entire TRN within each patient sample by treating the regulons as discrete units within the network (*i*.*e*., quantization) and calculating their individual activation status. For each regulon, we classified its status as over-expressed, under-expressed, or neither using a *p-*value cutoff of 0.05 for each patient sample as described in the Methods. Briefly, for a given regulon and patient sample, the number of genes in that regulon that appear in the top third of most highly expressed genes in that patient sample is compared to the number of genes expected to fall in the top third by random chance and a *p-* value is calculated. If the *p*-value is less than a threshold (e.g., 0.05), then the regulon is called over-expressed for that patient sample. Likewise, the number of regulon genes in the bottom third is used to determine if the regulon is under-active. A heatmap of the regulon activities across all patient samples reveals distinct patterns (**Fig. 2A**).

**Figure 2.**
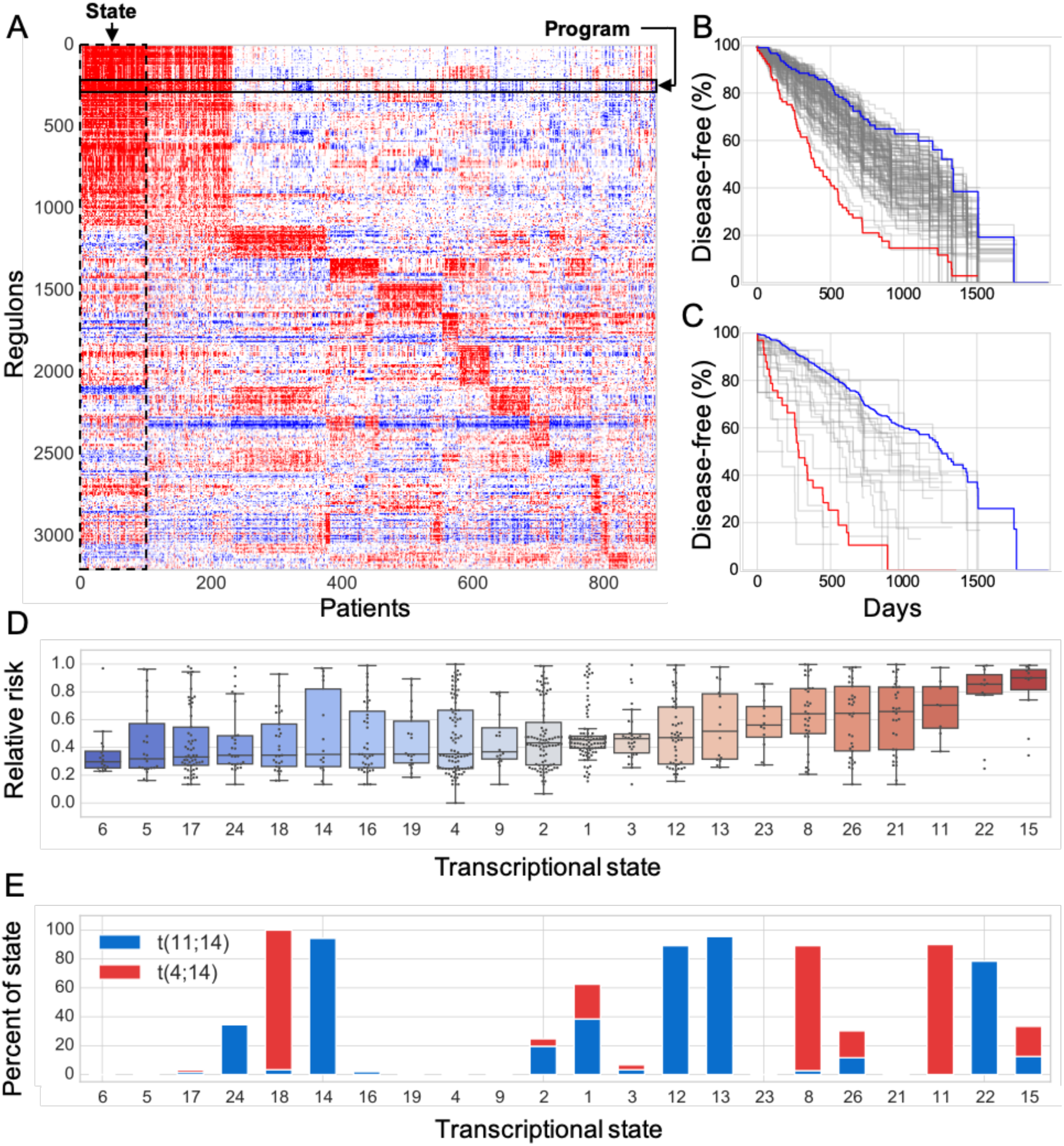
Programs and states. **A**. Heatmap of regulon activity across patients reveals distinct transcriptional states wherein patients have very similar regulon activity across the entire network. **B-C**. Kaplan-Meier survival curves showing the relationship between over-expression of each program (B) and state (C) and the observed progression-free survival in the CoMMpass study. Some programs and states are particularly high-risk (red) or low-risk (blue). **D**. The risk distribution of each state according to the Guan rank of the survival data. States with fewer than 5 patients represented in the survival data were omitted. **E**. Patients with t(4;14) or t(11;14) fall into several distinct transcriptional states with varying risk of disease progression.

### MM gene expression exhibits hierarchy of genetic programs and transcriptional states

Clusters of regulons, called *genetic programs* (or simply *programs*) herein, were observed to have similar activity across patient samples (Fig. 2A). This was often, but not exclusively, observed when multiple regulons had significant overlap of the genes that they contained. We note that different regulons can exhibit high degrees of gene overlap but have different associated regulators. Genetic programs are, therefore, a convenient way to uncover putative combinatorial regulation. In total, we discovered 141 programs, with an average of 90 unique genes and 21 unique transcription factors (i.e., 21 distinct regulons) per program. We evaluated the coherence of these genetic programs in two independent test datasets of MM (GSE24080^26^ and GSE19784^27^) by comparing the variance of the genes in each program against that of random selections of the same number of genes (500 permutations). 94.3% of the programs were coherent (variance < random, *p<*0.05) in GSE24080, and 92.2% were coherent in GSE19784, despite the fact that the training data (MMRF IA12) was collected via RNA-seq and the validation was microarray-based. The high degree of program coherence in multiple test datasets validates the generality of genetic programs discovered by our approach.

Regulon activity was also observed to cluster patients into subtypes of similar overall network activity profiles, which we call *transcriptional states* (**Fig. 2A**). Using our default clustering algorithm (see Methods), we discovered 25 distinct transcriptional states that accounted for 95% of the total patient population. The remaining 5% of patients did not match any of the states sufficiently well. We tested alternative clustering algorithms and found similar results. Interestingly, although some states are enriched with particular mutations, only t(4;14) and t(11;14) account for at least 90% of the samples within a single state. However, patients with these translocations are dispersed across several states, such that the presence of either translocation alone does not determine the particular transcriptional state observed. Thus, knowledge of individual mutations or chromosomal abnormalities is insufficient to predict the transcriptional state. Moreover, several alternative combinations of mutations can result in the same transcriptional state. For example, several states comprise some patients with t(4;14) and others with t(11;14).

### The CM TRN enables robust risk stratification

We performed univariate tests of all network feature types – mutations, programs, transcriptional states, regulon activity, and gene expression—to qualitatively evaluate the propensity of each to predict risk of disease progression (see Methods). The empirical distribution of uncorrected p-values from these univariate tests should be enriched in low values (i.e., *p* < 0.1) and relatively uniform (i.e., *p* >= 0.1) otherwise if the type of feature is a strong predictor. More formally, such a *p*-value distribution is indicative of a predictor that will have a lower false discovery rate than a predictor whose features are less enriched in the lower values.^28,29^ All feature types except mutations exhibited *p*-value distributions enriched in low values. Strong enrichments were observed for programs (58.9% satisfy *p* < 0.1), regulon activity (52.5% satisfy *p* < 0.1), and gene expression (46.8% satisfy *p* < 0.1), but not mutations (12.4% *P* < 0.1) (**Fig. S2**). It is especially noteworthy that the regulon activity distribution shows even greater enrichment than the gene expression distribution because the dimensionality has been significantly reduced and the values are discrete (i.e., activity = −1, 0, or 1). Aggregating the features in this way preserves and ostensibly enhances their ability to stratify risk. This supports the hypothesis that using regulons overcomes limitations of technical noise in gene expression measurements by averaging over large numbers of genes that are mechanistically co-regulated.

We tested the ability of a patient’s network status (*i*.*e*., the list of which regulons are activated and deactivated) to predict risk of disease progression using Ridge regression trained on the regulon activities of MMRF IA12. The predictive performance was evaluated in two microarray-based validation datasets of MM (GSE24080 and GSE19784) by first calculating the regulon activities in those datasets and then applying the predictor. Very strong performance was observed in all three datasets (**Table 1**), with validation AUC (*i*.*e*., area under the receiver operating characteristic curve) values of 0.70 in GSE24080 and 0.71 in GSE19784. This performance is on par with the best predictors available for MM and has the benefit of mechanisms and upstream causes associated to the predictive features. Thus, the status of the CM TRN can be inferred in individual patients and is predictive of risk for disease progression.

**Table 1.** Results of Ridge regression using regulon activity as features. N = number of samples, Median in days, Spearman correlation is predicted risk score vs GuanRank (i.e., observed risk rank), Cox regression is with respect to predicted risk score. * MMRF IA12 was used for training.

| Dataset | N | Median PFS | AUC | Spearman R | Spearman p | Cox HR | Cox p |
| --- | --- | --- | --- | --- | --- | --- | --- |
| *MMRF IA12 | 769 | 631 | 0.86 | 0.59 | 7.2 E-72 | 19.1 | 5.2 E-81 |
| GSE24080 | 559 | 1288 | 0.70 | 0.26 | 5.4 E-10 | 7.4 | 1.1 E-13 |
| GSE19784 | 282 | 782 | 0.71 | 0.35 | 1.4 E-9 | 6.3 | 2.7 E-10 |

Next, we performed Cox proportional hazards regression on the individual programs (**Fig. 2B**) and states (**Fig. 2C**) to quantify the extent to which these features stratify risk. Both the programs and states exhibited high- and low-risk features, with the states being particularly powerful determinants of risk (**Fig. 2D**). These observations stand in contrast to the results of individual mutations, which were not as powerful for stratifying risk, due in part to how infrequently most mutations appear. In general, the risk of a patient was better predicted by other patients with the same transcriptional state than by other patients with the same mutation. This is especially clear in the example of translocations t(4;14) and t(11;14) which exhibit distinct high-risk and low-risk transcriptional states (**Fig. 2E**).

As a final test of the information contained in the CM TRN we transformed the gene expression data to *network activity* by applying a correction to the measured expression value of a gene based upon the expression levels of the other genes to which it is mechanistically connected in the CM TRN. We compared the predictive performance of a gene’s expression to its network activity in the high-risk clinical subtypes to test whether the network correction improved predictive power. **Figure 3A-B** shows that the network activity preserves the large-scale patterns present in the gene expression data and appears much less noisy. For each high-risk clinical subtype, we classified the 30% of patients that were highest risk by GuanRank^30^ as truly high risk and classified the other 70% as low-risk. We then randomly split the samples into a training and test set (*i*.*e*., for each high-risk subtype), identified the gene that best stratified risk in the training set, and quantified its predictive performance in the test set by the area under the ROC curve (AUC). We repeated the prediction 100 times with random patient selections to generate a distribution of AUC scores. For all high-risk subtypes, the network activity of a gene was a better predictor of risk than the gene expression data (**Fig. 3C**). The sub-stratification of progression-free survival enabled by the network activity of single genes is shown by subtype in **Figure 3D**. Finally, the genes that were most predictive of risk in MMRF IA12 across all subtypes were evaluated in the GSE24080 and GSE19784 test sets. In both cases, the network activity outperformed the gene expression for predicting risk (**Fig. S3**). The ability of the CM TRN to improve the predictive performance of individual genes and apparently filter noise from the corresponding expression data are strong indicators that the relationships in the inferred network reflect meaningful biological mechanisms.

**Figure 3.**
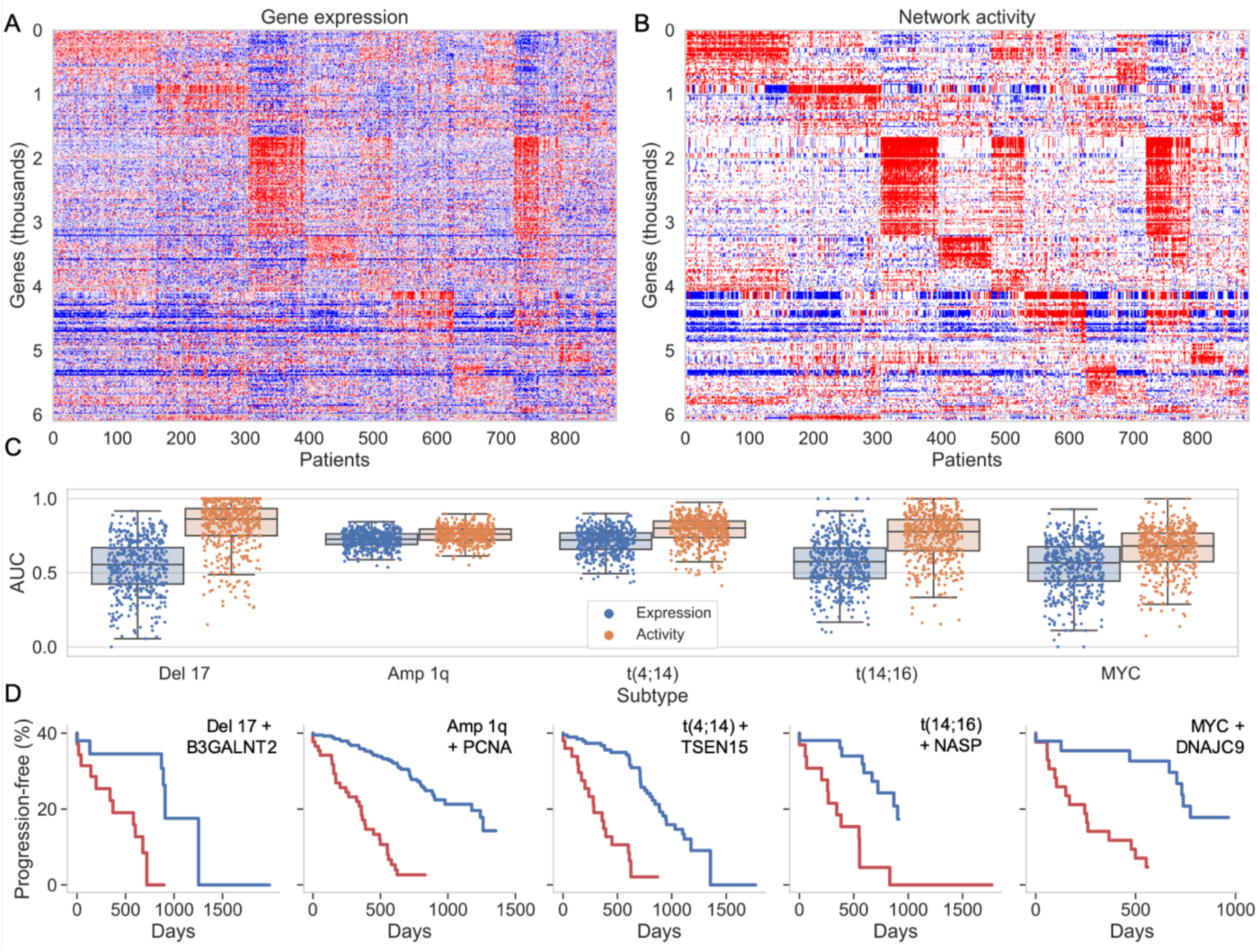
Network activity. **A-B**. Heatmaps of (A) normalized gene expression and (B) network activity for all genes appearing in at least two regulons. **C**. Boxplots comparing the LASSO prediction of high-risk subsets using normalized expression (blue) or network activity (yellow) of the genes as features. Each point overlaying the boxplots is a mean area under the ROC curve (AUC) from 5-fold cross-validation of the high-risk prediction. 100 iterations were performed for each condition to avoid bias in the selection of training and test subsets. **D**. Kaplan-Meier curves demonstrating the sub-stratification of clinical subtypes based upon the network activity of the most predictive individual or pair of genes as determined by the LASSO predictions in C.

### High-risk genetic program underlies proliferation and recapitulates predictive signatures

We analyzed the highest-risk genetic program, Pr-68 (Cox HR = 8.8, *p <* 1.3e-18), to better understand the mechanisms most strongly correlated to rate of disease progression. The genes in Pr-68 are heavily enriched in DNA-replication (*p* < 1e-25), cell cycle (*p* < 1e-25) and DNA mismatch repair (*p* < 1e-8) functions, and the regulons comprising Pr-68 are enriched for 10 hallmarks of cancer. Thus, Pr-68 is a genetic program associated with proliferation. The genes of Pr-68 include BRCA1, RAD51, RAD51AP, and PARPBP, and PARP1 is among its regulators. These genes reflect the previous report of “BRCAness” that is observed after escape from proteasome inhibitors and confers sensitivity to PARP inhibitors.^31^ Several studies have identified proliferation signatures as optimal risk classifiers, but the underlying biological drivers have not been clearly established. We find that the genes of Pr-68 have an adjusted p-value of 1e-55 for enrichment in ChIP-seq targets of FOXM1, suggesting that FOXM1 is an important upstream regulator and potential therapeutic target. We note that FOXM1 has been experimentally confirmed as an important target in high-risk MM.^32^ FOXM1 is most significantly activated by E2F1 in the MM TRN (Spearman correlation: *p < 1e-172; E2F1 motif in FOXM1 promoter*). All high-risk subtypes, with the exception of t(4;14), were causally upstream of FOXM1 activation via the intermediate up-regulation of E2F1 in the CM TRN.

Four existing prognostic gene expression profiles of MM demonstrated significant overlap with the genes of Pr-68. We compared the gene sets of UAMS70^33^, EMC92^34^, M3CN^24^, and the Proliferation signature of Hose *et al*.^35^ to the Pr-68 genes and computed hypergeometric *p-*values. The results, listed in **Table 2**, show that the overlap of all four signatures with Pr-68 were highly significant and cumulatively accounted for 108 of the 228 (47%) genes in Pr-68. Moreover, the gene PHF19, whose expression was identified as the best individual predictor of risk in a recent MM DREAM challenge^36^, is also a member of Pr-68.

**Table 2.** Existing prognostic signatures of high-risk MM map to Pr-68. All four of the prognostic signatures exhibit significant overlap with Pr-68. *Only 65/70 UAMS probes had gene names.

| Prognostic signature | Reference | Overlap with Pr-68 | p-value |
| --- | --- | --- | --- |
| UAMS70 | Shaughnessy Jr, *et al.* | 8 / 65* | $3.0 \times 10^{-4}$ |
| EMC92 | Kuiper, *et al.* | 17 / 92 | $2.2 \times 10^{-10}$ |
| Proliferation | Hose, *et al.* | 40 / 60 | $2.7 \times 10^{-55}$ |
| M3CN | Liu, *et al.* | 82 / 178 | $1.2 \times 10^{-85}$ |
| PHF19 | Mason, *et al.* | 1 / 1 | $2.7 \times 10^{-2}$ |

Several existing therapies map to the genes of Pr-68, and thus may be good candidates for treating high-risk MM. We searched the OpenTargets database for MM therapies that target the genes comprising Pr-68 and their regulators. In total, 26 drugs mapped to Pr-68. We note that FOXM1 and E2F1 appear to be master regulators of Pr-68, given the number of genes that they regulate within the program, but we did not find any existing therapies that had been approved for them.

### CRBN activity is linked to high-risk genetic program through CCNDBP1

Activation of E2F1 and FOXM1, and thus the genes of Pr-68, is known to occur via the CCND1-CDK4 complex.^37–39^ The cyclin D binding protein CCNDBP1 has previously been shown to interfere with the CCND1-CDK4 complex, providing a putative mechanism to prevent Pr-68 activation and thereby halt the G1/S transition of the cell cycle.^40,41^ We find that CCNDBP1 is strongly deactivated in canonical MM translocation subtypes relative to other subtypes (p < 1 x 10^−73^ via Wilcoxon rank-sum test). This suggests that the translocations confer greater dysregulation of CCND1, facilitating constitutive activation of the cell cycle.

Inspection of the CM TRN reveals that CCNDBP1 belongs to a regulatory circuit that may provide critical insight for the treatment of MM. In particular, the transcription factors that drive activation of CCNDBP1 in the CM TRN are directly linked to the activation of the IMiD substrate CRBN (**Fig. S4**).^42^ The most effective mechanism to down regulate either CCNDBP1 or CRBN in the context of the CM TRN is deactivation of transcription factor ZNF35. This means that down-regulating CCNDBP1 to facilitate dysregulation of the CCND1-CDK4 complex has the consequence of down-regulating CRBN, and thus interfering with the IMiD therapeutic mechanism of action. Conversely, down-regulating CRBN to escape IMiD therapy has the consequence of down-regulating CCNDBP1, thus dysregulating the CCND1-CDK4 complex and facilitating greater proliferation. Indeed, the network activity of CRBN and CCNDBP1 are strongly correlated (Spearman correlation: R = 0.62, *p < 1 x 10*^*-94*^), as is their gene expression (*i*.*e*., without network correction) (Spearman correlation: R = 0.51, *p < 1 x 10*^*-60*^). Moreover, the activity of CRBN, like CCNDBP1, is significantly lower in translocation subtypes (p < *7 x 10*^*-60*^ via Wilcoxon rank-sum test).

### Drug targets show subtype-specific risk stratification

We searched for subtype-specific relationships to risk of disease progression in both the network activity and standard gene expression of genes whose corresponding protein is targeted by therapies available for MM. We considered both network activity and gene expression because they have complementary strengths. In theory, a gene that is well connected in the network will benefit from this correction, but a gene that is poorly connected (e.g., present in only 1 regulon) may be over-corrected due to a lack of information, such that the measured gene expression is more reliable. The correlation between risk of disease progression to network activity and gene expression is provided for each drug target by subtype in **Tables S1-S2**. The network activity of these important targets is visualized by risk decile for each subtype in **Figure 4**. Several noteworthy patterns emerge.

**Figure 4.**
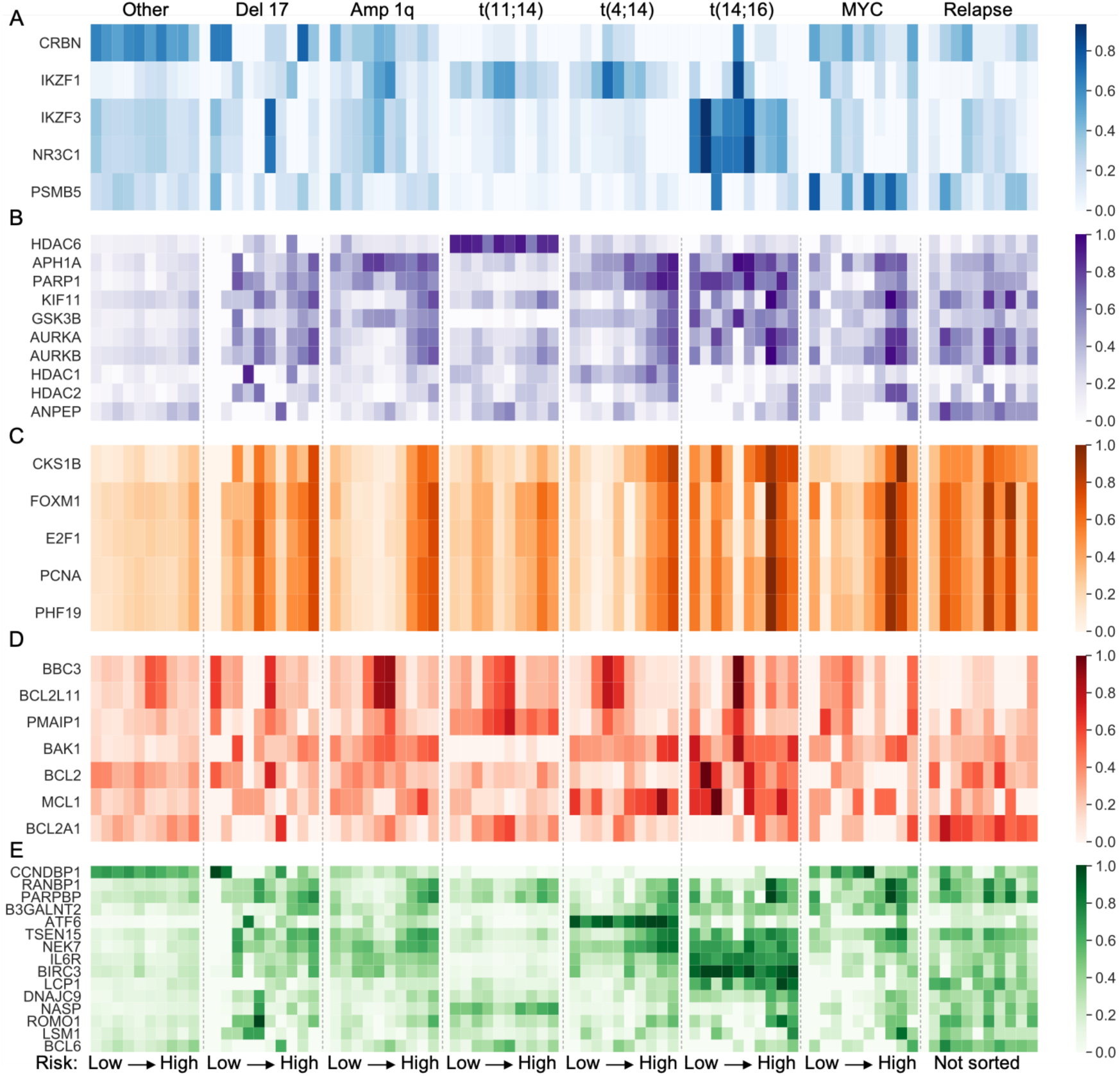
Network activity of drug targets across subtypes. **A**. Targets of established baseline therapies. IMiDs: CRBN, IKZF1, IKZF3; Dexamethasone: NR3C1; Bortezomib and Carfilzomib: PSMB5. **B**. Targets of therapies in clinical trials for relapse-refractory MM (RRMM). **C**. Genes associated with proliferation. **D**. Genes directly involved in the intrinsic apoptosis pathway demonstrate subtype-specific differences in network activity. **E**. Risk-associated and novel target genes. These genes either classify a subtype via their uniformly high expression or are selectively active in the low- or high-risk subset of a clinical subtype.

### IMiDs – CRBN, IKZF1, IKZF3

All three IMiD targets^42^ exhibited some degree of subtype-specific risk stratification. In addition to the significant difference in activity between translocation and non-translocation-bearing patients, CRBN sub-stratified risk in t(4;14) (*p*_*expression*_*<1*.*5 x 10*^*-3*^, *p*_*activity*_*<0*.*1*), Amp 1q (*p*_*expression*_*<6*.*1 x 10*^*-2*^, *p*_*activity*_*<4*.*6 x 10*^*-2*^), and patients without high-risk features or t(11;14) (*p*_*expression*_*<7*.*9 x 10*^*-2*^, *p*_*activity*_*<3*.*1 x 10*^*-2*^, labeled “Other” in **Tables S3-S4**). In each case, the CRBN activity was anticorrelated to risk, such that high activity of CRBN was favorable. This is in agreement with the role of CRBN in the mechanism of IMiD therapy, wherein it acts as a substrate for the degradation of IKZF1 and IKZF3. IKZF1 expression did not demonstrate significant correlation to risk in any of the subtypes, but IKZF1 network activity was significantly anticorrelated with risk in t(4;14) patients (*p*_*activity*_*<3*.*5 x 10*^*-2*^). The low expression levels of IKZF1 and high coverage in the network (*i*.*e*., member of 14 regulons) are precisely the conditions in which we would expect network activity to outperform standard gene expression. The higher activity of IKZF1 in low-risk t(4;14) patients is in agreement with its importance as a degradation target in the CRBN-dependent mechanism of IMiD therapy. IKZF3 gene expression was not correlated to risk in any subtypes, but IKZF3 network activity was anticorrelated to risk in t(14;16) patients (*p*_*activity*_*<2*.*1 x 10*^*-2*^). This may indicate a therapeutic benefit to the IMiD-based degradation of IKZF3 in t(14;16) patients.

### Dexamethasone – NR3C1

The gene expression and network activity of the dexamethasone target^43^ *NR3C1* were both significantly anticorrelated to risk in t(14;16) patients (*p*_*expression*_*<3*.*4 x 10*^*-4*^, *p*_*activity*_*<1*.*5 x 10*^*-2*^). The strong association between low levels of NR3C1 and high risk in t(14;16) indicate that escape from dexamethasone therapy is particularly important in this subtype. It is unclear, however, if the mechanism of escape involves down-regulation of NR3C1 within MM cells, or a smaller proportion of NR3C1-expressing cells (*e*.*g*., in the microenvironment). The activity profile of NR3C1 is very similar to IKZF3, as these genes have similar connectivity in the CM TRN. Specifically, they are both strongly activated by ZEB1 and RORA. Thus, activation of these TFs may stabilize the high target activity of IKZF3 and NR3C1.

### Bortezomib – PSMB5

The functional target of frontline proteasome inhibitors, PSMB5^44^, shows relatively little subtype-specific correlation to risk at baseline. Although the gene expression of PSMB5 did not significantly correlate to risk in any subtype, its network activity was significantly correlated to risk in t(4;14) (*p*_*activity*<_0.05). In particular, patients with low PSMB5 activity (*i*.*e*., PSMB5_activity_<0) tended to exhibit lower risk than patients with moderate to high activity (*p<*7.1 x 10^−2^).

### HDACs – HDAC2 and HDAC6

HDAC expression profiles vary dramatically between subtypes. In particular, HDAC6 is ubiquitously expressed in t(11;14) patients, but not in other subtypes. Because HDAC6 is highly expressed across t(11;14) patients it does not correlate to their risk, however it may serve as a promising target in these patients if its expression reflects an essential function played in t(11;14) MM cells. Among the other HDACs, HDAC2 stood out for its subtype-specific correlation to risk, particularly in t(4;14) (*p*_*expression*_*<5*.*6 x 10*^*-3*^, *p*_*activity*_*<7*.*7 x 10*^*-4*^). HDAC2 also correlated to risk in Amp 1q (*p*_*expression*_*<8*.*1 x 10*^*-2*^, *p*_*activity*_*<4*.*6 x 10*^*-2*^) and t(14;16) (*p*_*expression*_*<6*.*2 x 10*^*-2*^, *p*_*activity*_*<6*.*8 x 10*^*-2*^) patients, though not as strongly.

### Mitosis-related targets – AURKA, AURKB, KIF11

The mitosis-related targets AURKA, AURKB, and KIF11 are all members of Pr-68 and stratify risk well across several subtypes. AURKA, AURKB, and KIF11 are significantly correlated to risk in all subtypes other than t(14;16) and MYC (*i*.*e*., patients with MYC overexpression). Although the network activity of KIF11 does not significantly correlated to risk in the t(14;16) and MYC subtypes, its gene expression does. The most significant risk stratification exhibited by these genes occurs in t(4;14) patients (AURKA: *p*_*expression*_*<2*.*2 x 10*^*-5*^, *p*_*activity*_*<5*.*7 x 10*^*-6*^, AURKB: *p*_*expression*_*<1*.*6 x 10*^*-4*^, *p*_*activity*_*<1*.*6 x 10*^*-5*^, KIF11: *p*_*expression*_*<7*.*2 x 10*^*-5*^, *p*_*activity*_*<1*.*0 x 10*^*-4*^)

### PARP inhibitors – PARP1

The network activity of PARP1 correlated to risk in all subtypes other than t(11;14), t(14;16), and MYC. The correlation to risk was greatest in t(4;14) patients (*p*_*expression*_*<6*.*8 x 10*^*-4*^, *p*_*activity*_*<3*.*5 x 10*^*-6*^). We note that our CM TRN shows that PARP1 is an activator of TOP2A and NEK2. Inhibition of PARP1, TOP2A, or NEK2 have all independently been shown to overcome resistance to proteasome inhibitors.^31,45,46^ The high activity of these genes in the highest-risk t(4;14) patients at baseline suggests that these patients would quickly become resistant to proteasome inhibitors, but would benefit from PARP1 inhibitors. We note that pegylated lysosomal doxorubicin targets TOP2A^47^ and could also benefit high-risk t(4;14) patients.

### Monoclonal antibodies – APH1A (gamma-secretase)

Monoclonal antibodies have emerged as promising next-generation therapies for MM but are susceptible to resistance by over-activation of the gamma-secretase protein complex, which cleaves the antibody targets from the cells. APH1A is an essential component of this complex and exhibits subtype-specific risk correlation. The APH1A gene is located on chromosome 1q and exhibits high network activity across the Amp 1q subtype, such that Amp 1q patients may be especially prone to gamma-secretase-mediated resistance to mAbs. Similar to PARP1, the network activity of APH1A correlated to risk in all subtypes other than t(11;14), t(14;16), and MYC. This motivates therapy by gamma-secretase inhibitors in these patients regardless of monoclonal antibody use, but especially in conjunction with them. We note that APH1A expression strongly correlates to PARP1 (*p*<*3*.*0 x 10*^*-31*^), TOP2A (*p*<*3*.*6 x 10*^*-6*^), and NEK2 (*p*<*6*.*0 x 10*^*-19*^), so it is unsurprising that the t(4;14) patients exhibit the strongest risk stratification by APH1A activity (*p*_*expression*_*<1*.*2 x 10*^*-2*^, *p*_*activity*_*<4*.*8 x 10*^*-5*^).

### BCL2-family – BCL2, MCL1, NOXA, BAK1, BCL2A1

Figure 4D shows subtype-specific activity profiles of BCL2-related genes involved in the intrinsic apoptosis pathway. Patients harboring translocation t(11;14) exhibit a unique profile of these genes (**Fig. S5**), with high PMAIP1 (also called NOXA) and low BAK1 activity, such that these patients can be classified by their high PMAIP1/BAK1 ratio. It has been reported that the ability of PMAIP1 to bind the anti-apoptotic protein MCL-1 may underlie the success of BCL2-inhibitors in some patients.^48^ Our network analysis supports that claim insofar as PMAIP1 shows much higher activity in t(11;14) than in other subtypes, and the responders to BCL2-inhibitors in MM have been significantly enriched with t(11;14) patients.^49^ We note that the BCL2-family protein BCL2A1 is an interesting novel target for RRMM, as it becomes significantly over-expressed at relapse (*p*<*2*.*7 x 10*^*-5*^).

### Melflufen – ANPEP

The Melphalan-flufenamide conjugate, Melflufen, targets MM cells that over-express the protein product of ANPEP and has shown clinical benefit in RRMM patients. We find that the network activity and gene expression of ANPEP are both mildly correlated to risk in patients that do not bear high-risk features or t(11;14) (*p*_*expression*_*<1*.*4 x 10*^*-2*^, *p*_*activity*_*<1*.*7 x 10*^*-2*^). ANPEP network activity, but not gene expression, is also correlated to risk in t(14;16) patients (*p*_*activity*_*<2*.*3 x 10*^*-3*^). The most noteworthy trend in ANPEP expression, however, is that it becomes more highly expressed at relapse (*p<7 x 10*^*-3*^), in accordance with the success of Melflufen in RRMM.^50^

### Population at relapse

There are only 39 patients with baseline gene expression, first relapse gene expression, and clinical outcomes data in MMRF IA12. Thus, pairwise comparisons are feasible, but limited in sample size. Of these 39 patients, 21 had no translocations or high-risk features at baseline, 8 were t(11;14) subtype, 7 exhibited Amp 1q, 4 were t(4;14) subtype, 2 exhibited MYC overexpression, and 1 exhibited Del 17. Therefore, pairwise comparisons by subtype are statistically underpowered, with the possible exception of patients with no translocations or other high-risk features. Moreover, these 39 patients were significantly higher risk (*p*<3 x 10^−7^) than the remaining baseline patients that did not have matched relapse profiles in MMRF IA12, presumably because these patients relapsed faster and thus were the ones with data available. Given the limited relapse data available, we pooled all relapse samples (N=56) with expression, even though 17 samples did not have matching baseline expression and clinical outcomes, and compared these profiles against the pool of all baseline expression profiles.

### Differential activity of drug targets at relapse

Both network activity and gene expression showed decreased activity of CRBN at relapse when pooling all baseline versus all first-relapse patients (*p*_*expression*_*<1*.*5 x 10*^*-3*^, *p*_*activity*_*<1*.*1 x 10*^*-2*^). However, only network activity showed a significant difference (*p<*0.05) in the pairwise comparison of CRBN in the 39 patients with matched data. The network activity of IKZF1 is also significantly higher at baseline than relapse (*p*_*activity*_<4.3 x 10^−4^), but the expression of IKZF1 alone does not show significant difference (*p*_expression_>0.8). Decreased CRBN activity at relapse suggests a possible mechanism of escape from IMiD therapy by limiting the substrate for IKZF1 degradation, whereas decreased IKZF1 activity at relapse suggests a possible mechanism of escape wherein the MM cells are no longer dependent upon the activity of IKZF1. The expression of CRBN and IKZF1 are well correlated at relapse (spearman R=0.40, *p<2*.*3 x 10*^*-3*^), suggesting that the two possible mechanisms are not occurring in distinct subsets of relapse patients (*i*.*e*., IKZF1 is not lowly expressed when CRBN is highly expressed).

### Relapse is characterized by differential activity of 5 genetic programs

The genes with greatest differences in network activity between baseline and relapse largely fall into 5 programs: Pr-0, Pr-4, Pr-34, Pr-68, and Pr-134 (**Fig. 5**). The program with the most extreme deactivation at relapse is Pr-34, which notably contains the genes IKZF1 and PSMB7. Program Pr-0 is also strongly deactivated at relapse, and notably contains several pro-apoptotic genes: BCL2L11 (BIM), BBC3 (PUMA), BCL7B, TP53BP2, and TP53INP2. On the other hand, programs Pr-68 and Pr-134 are the most strongly activated at relapse. Pr-68 is the aforementioned genetic program driven by FOXM1 that characterizes high risk at baseline and is enriched with markers of proliferation. Interestingly, Pr-134 is not associated with risk at baseline (Cox HR = 1.0, *p>0*.*31*).

**Figure 5.**
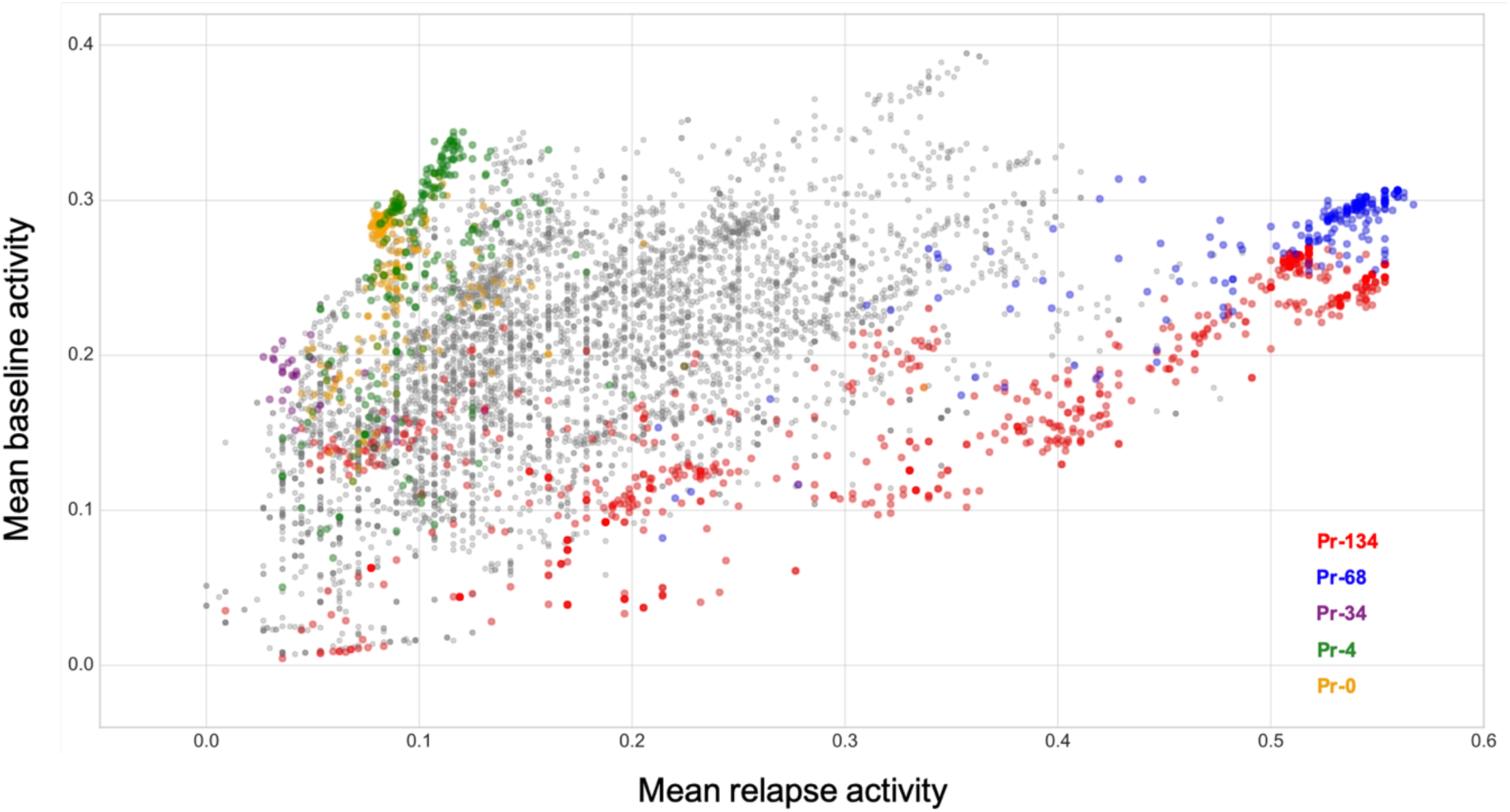
Network activity of genes at baseline versus relapse. Each point depicts the mean network activity of a gene at relapse versus baseline. Many genes have significantly different network activity at relapse versus baseline, with the most extreme differences observed for the genes of programs Pr-134, Pr-68, Pr-34, Pr-4, and Pr-0.

**Figure 6.**
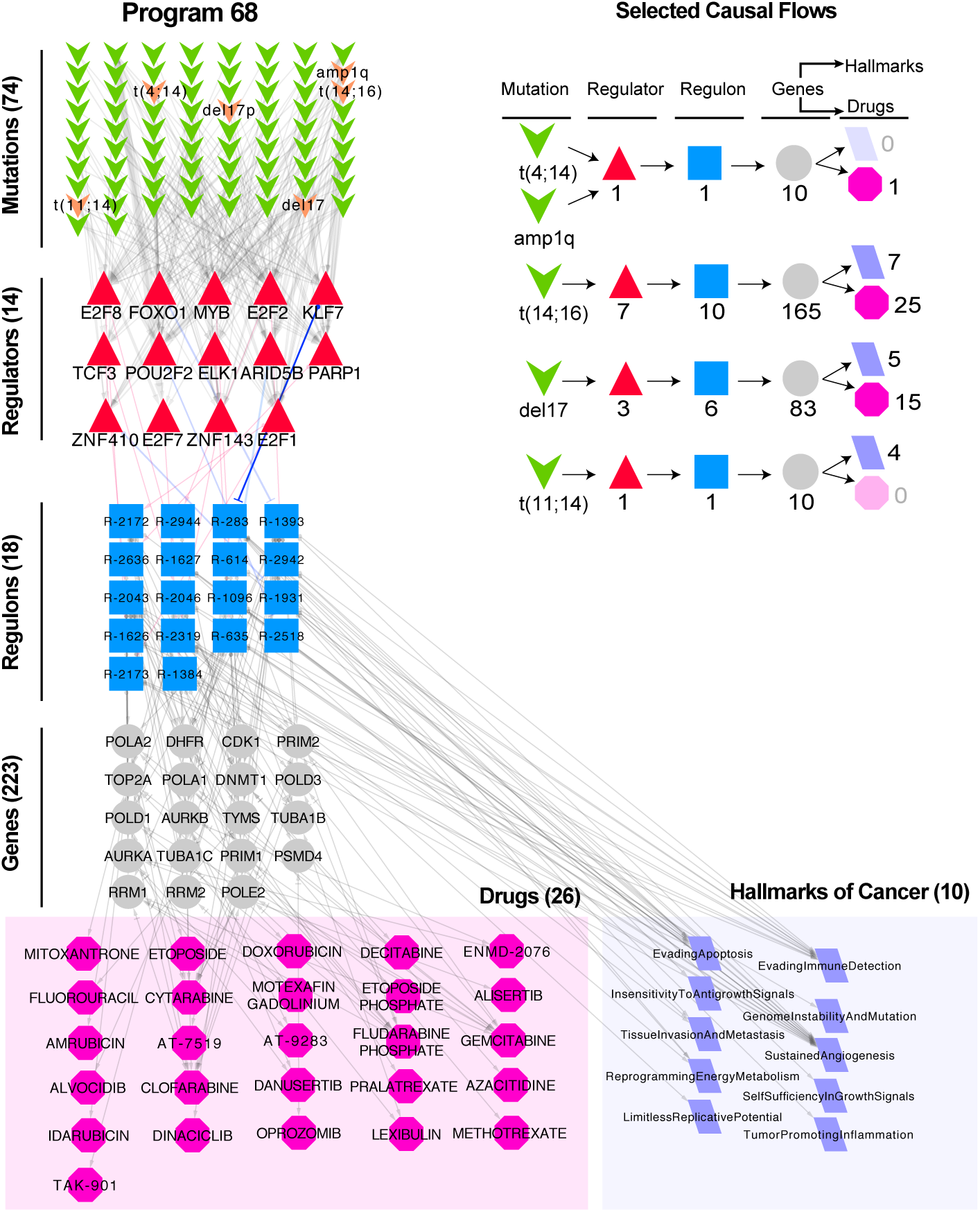
Network visualization of Pr-68. The regulons comprising program Pr-68 are represented as blue boxes with their associated regulators (red triangles) and putative causal genetic abnormalities (green chevrons). Hallmarks of cancer that are associated with these regulons are highlighted by purple parallelograms, and drugs that target genes (gray) within these regulons are highlighted by pink octagons.

### The genetic program with highest activity at relapse reflects the microenvironment

Whereas Pr-68 reflects mechanisms of proliferation, Pr-134 comprises many markers of the immune-suppressed microenvironment. Especially noteworthy are signatures of myeloid-derived suppressor cells (MDSCs). In addition to the characteristic surface marker CD11b, many of the cytokines that promote the recruitment or generation of MDSCs (e.g., M-CSF, G-CSF, IL-18, IL-1B, IL-10, CCL2, S100A8, S100A9, PTGER4)^51^ are present in Pr-134. These cytokines, and those produced by the MDSCs themselves, are known to promote an immunosuppressive microenvironment. Signatures of bone marrow stromal cells (BMSCs), M2-polarized macrophages, mesenchymal stem cells (MSCs), osteoclasts, non-cytotoxic T cells, anergic exhausted cytotoxic T cells, NK cells, and cancer-associated fibroblasts (CAFs) are also present in Pr-134 (**Fig. 7**).

**Figure 7.**
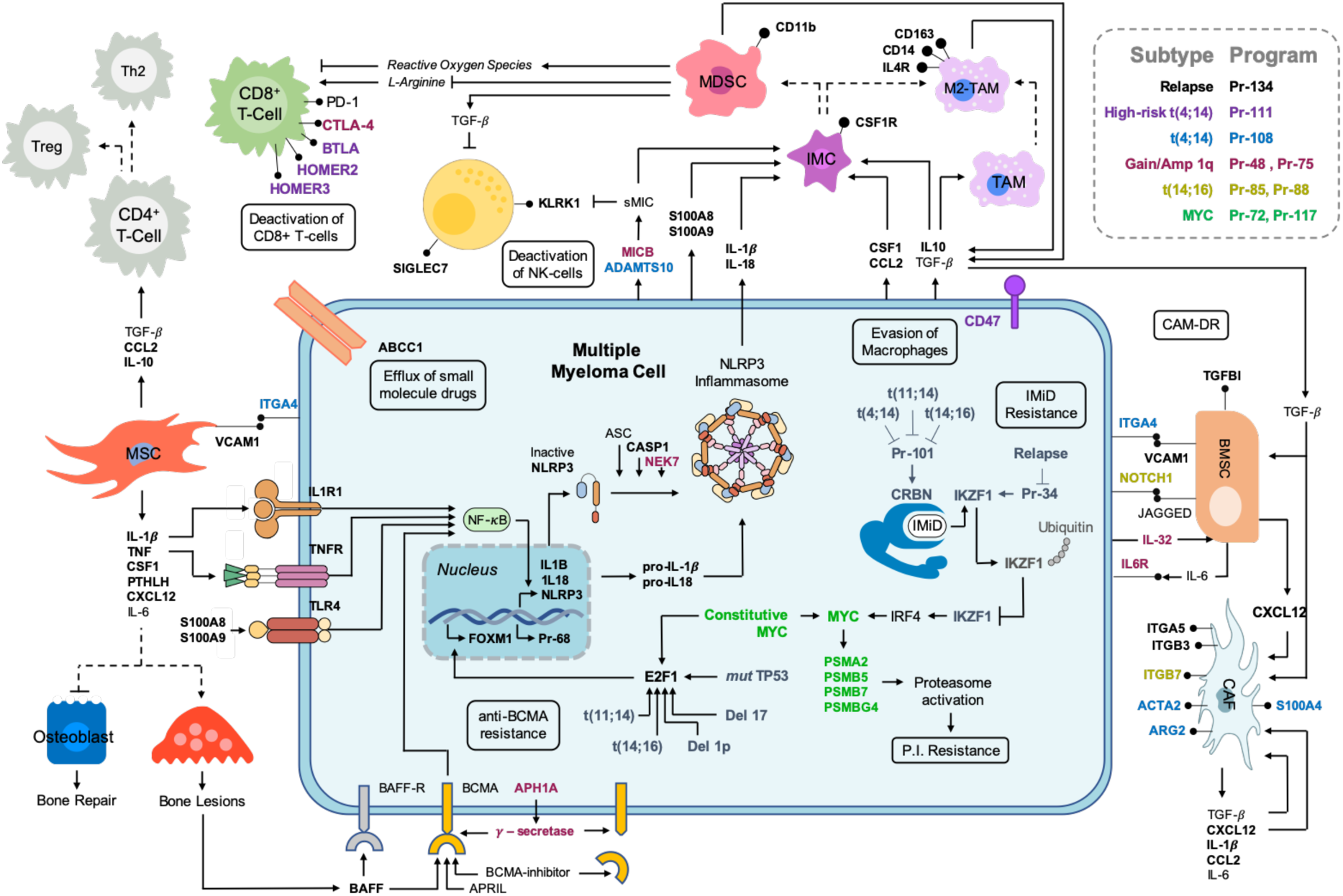
Pr-134 schematic. The genes of Pr-134 include markers of cell types characteristic to the bone marrow (BM) microenvironment, NF-kB signaling, and the NLRP3 inflammasome. The high activity of these genes at relapse suggests the engagement of the BM microenvironment to facilitate escape from therapy and evasion of immune cell-mediated killing. The high-risk clinical subtypes are complementary to Pr-134 in escaping therapy and immune surveillance and they promote proliferation of multiple myeloma cells by activating Pr-68 via E2F1 and FOXM1. Bold gene names indicate that the gene is present in the program of the corresponding color (see legend).

## Discussion

Multiple myeloma has been shown to present heterogeneity of cytogenetics, mutations, gene expression, and clinical outcomes. However, we lack a comprehensive map of the mechanistic links between these features. In particular, a more complete picture of the molecular signatures that are dysregulated across disease subtypes is necessary to understand the downstream effects of mutations and the points of actionable intervention in their mechanisms. We developed MINER to uncover the structure and hierarchy of patterns in gene expression from a mechanistic and causal perspective in MM. Despite the myriad combinations of mutations and chromosomal aberrations observed in MM patients, the transcriptional data is relatively well structured into 25 transcriptional profiles with 141 programs that can be further divided into 3,203 regulons. This highlights a critical opportunity in the systems biology of cancer: The effect of the myriad possible combinations of mutations is too great to study with statistical power in clinical analysis because specific combinations are too rare, but the end-result is a discrete set of transcriptional profiles that *can* be studied effectively. Indeed, we see that transcriptional features such as genetic programs, regulons, network-constrained gene activity (*i*.*e*., network activity), and transcriptional states stratify risk of disease progression better than mutations in the case of MM (**Fig. S2**). Studying the transcriptional landscape of a disease in the context of a CM TRN enables relationships between the activities of genes and clinical outcomes to be traced to the putative causal effects of mutations. Moreover, those mutations that are known to predict risk can be better understood by investigating their downstream effects in the CM TRN.

Diverse biological mechanisms including variable expression of drug targets, distinct profiles of apoptosis regulators, or differing degrees of cell proliferation at baseline could conceivably underly the risk of disease progression. Comparison of these effects in the context of the CM TRN showed that individual genes (*e*.*g*., CKS1B, PCNA, E2F1, FOXM1, PHF19) and genetic programs (*e*.*g*., Pr-68) associated with cell proliferation were the best predictors of risk. This agrees with previous studies but doesn’t address the root cause of the cellular proliferation. However, we can use the CM TRN to see which mutations and regulators are upstream of these proliferative signatures in each clinical subtype of MM. We find that all high-risk clinical subtypes except t(4;14) are causally linked to the activation of Pr-68 by promoting activation of FOXM1 via up-regulation of E2F1. We hypothesize that FOXM1 is a master regulator of Pr-68 given that the genes in Pr-68 are highly enriched with confirmed ChIP-seq targets of FOXM1 (*p<*1.0 x 10^−55^). Activation of E2F1 occurs as a direct consequence of dysregulating the G1/S cell cycle checkpoint via the common chromosomal aberrations observed in MM.

Although proliferation signatures are most predictive, the activity of drug targets within the CM TRN also stratify risk at baseline. This is especially noteworthy in the case of the IMiD substrate CRBN, which is highly active in the subset of patients without clinical high-risk features. These patients do not exhibit strong proliferation signatures at baseline, so they likely benefit from low baseline aggressiveness of disease and high susceptibility to IMiD therapy. Subtype-specific differences in network activity suggest that complimentary MM treatments can be particularly effective for specific subsets of MM patients. In particular, HDAC6 inhibitors are rational therapies for t(11;14) patients, as they ubiquitously express HDAC6. Although experimental confirmation is required, this may signal a critical dependency of t(11;14)-driven MM on HDAC6. Moreover, the especially high activity of PMAIP1 in t(11;14) supports the observed benefit of BCL2-inhibitors in this subtype. Translocation t(4;14) also exhibits actionable subtype-specific risk signatures. The activity of PARP1 and TOP2A each correlate with high risk in t(4;14) and enable proteasome resistance. Thus PARP-inhibitors and liposomal doxorubicin (*i*.*e*., targeting TOP2A) are rational therapies for t(4;14) patients that become resistant to proteasome inhibitors. Additionally, the correlation of APH1A activity to risk in t(4;14) and Amp 1q patients suggests that a gamma-secretase inhibitor may be particularly important for regimens involving mAbs. This agrees with the observation that gain of 1q21 confers poor prognosis in patients treated with Daratumamab (*i*.*e*., anti-CD38 mAb).^52^ Finally, we note that virtually all subtypes exhibit correlation between risk and the activity of the mitosis-related targets AURKA, AURKB, and KIF11, which may indicate that these therapies will be effective in high-risk MM.

While the profile of apoptosis regulators was not strongly predictive of risk at baseline, significant changes were observed at relapse. Program Pr-0, which includes several pro-apoptotic regulators, shows much lower activity at relapse. Although BCL2 and MCL1 did not exhibit noteworthy changes at relapse, BCL2A1 became ubiquitously active. Accordingly, we anticipate that BCL2A1 may be an important novel target in RRMM.

A surprising result of this research was the observation of a strong signature of the immune microenvironment, despite the RNA-seq data being obtained from CD138+-purified MM cells. Markers of virtually all cells previously implicated in BM microenvironment-mediated resistance to therapy are present in program Pr-134, which becomes highly activated at relapse. This provides support to the hypothesis of a microenvironment-driven mechanism of relapse to therapy. The observation of these signatures in purified MM cells suggests that the mechanism of resistance may involve exosomes or other methods of delivering RNA from microenvironment cells. Although it is not yet clear which therapeutic strategies will be most effective for interfering with microenvironment-driven resistance, we note that targets related to NF-kB signaling may be especially important. In particular, activation of NF-kB appears to be central to communication between the microenvironment and MM cells in the context of the CM TRN. NF-kB signaling is known to activate the NLRP3 inflammasome which generates IL-18 and IL-1B.^53^ These ligands stimulate cells of the microenvironment, such as MDSCs^54^, to ultimately produce ligands such as S100A8, S100A9, TNF, and IL-1B, all of which stimulate receptor-mediated NF-kB signaling and thus complete a circuit.^51,53,55–58^ The genes of this circuit belong to Pr-134, suggesting that NF-kB signaling is a driver of microenvironment-induced resistance.

Although Pr-134 is not associated with risk at baseline, other signatures of interaction with the immune microenvironment are observed in high-risk MM patients. For example, program Pr-108, activated across all t(4;14) patients, contains ITGA4, which can directly engage BMSCs via VCAM1.^59^ Moreover, the highest-risk subset of t(4;14) patients over-activate program Pr-111, which contains the “don’t eat me” marker CD47 that evades killing by macrophages^60,61,62,63^, and BTLA, HOMER2 and HOMER3, all of which suggest escape from cytotoxic T-cell killing. Patients with amp 1q exhibit overactivation of Pr-52 and Pr-75, which express genes that directly promote the differentiation of osteoclast progenitor cells into osteoclasts (e.g., ANXA2).^64^ Moreover, Pr-52 and Pr-75 contain MICB, which in conjunction with ADAMTS10 contained in t(4;14)-activated program Pr-108, generates soluble MICB (sMICB).^65^ Soluble MICB has been shown to inactivate NK cells and cytotoxic T cells, steer macrophages to the tumor-promoting M2 phenotype, and stimulate the generation of MDSCs.^66–69^ Lastly, Pr-75 contains IL6R and IL32, which promotes osteoclastogenesis and stimulates BMSCs to produce IL6.^70,71^ Taken together, the “double-hit” combination of amp 1q and t(4;14) bears signatures of a microenvironment that inactivates killing by cytotoxic lymphocytes, promotes formation of immuno-suppressive MDSCs and M2-polarized macrophages, and activates paracrine IL6 signaling. We note that the presence of both t(4;14) and amp 1q show synergistic activation of Pr-75 and Pr-134, such that patients harboring both abnormalities over-activate Pr-75 (*p<5*.*1 x 10*^*-8*^) and Pr-134 (*p<4*.*9 x 10*^*-7*^) relative to patients with either amp 1q or t(4;14) alone. This suggests an immune-suppressive synergy as one element of the elevated risk of t(4;14)-amp 1q “double-hit” patients.

## Conclusion

The coherence of inferred programs across datasets, strong performance of network-activity based predictions, and recapitulation of therapeutic escape mechanisms all provide support for the inferred network and methods of analysis reported herein. Although the transcriptional profiles of MM patients are heterogeneous, they are well-described by 141 genetic programs and 25 transcriptional states of differing risk for disease progression. Changes to the relative activities of these programs across MM subtypes offers actionable insights into the mechanisms of resistance or susceptibility to various therapies, with promising new insights for relapse refractory patients.

## Methods

### Data selection

In order to generate and test a MINER TRN of multiple myeloma, we identified and pre-processed multiple publicly-available datasets. Several gene expression datasets with associated clinical outcomes exist for multiple myeloma, but the most extensive is that provided by the Multiple Myeloma Research Foundation (MMRF) as a result of their CoMMpass study. In total, 1150 patients from 90 worldwide sites had bone marrow samples analyzed every 6 months for 8 years. The samples were subject to many types of analysis, including genomic, cytogenetic, and transcriptomic analysis via RNA-seq of CD138^+^-purified MM cells. However, not all data from the CoMMpass study was publicly available at the time of this work. Nonetheless, the 881 samples with RNA-seq and translocation calls, 769 samples with matched clinical outcomes, and 734 samples with somatic mutation data available in the MMRF interim analysis 12 (IA12) represents the richest multiple myeloma dataset at the time of our analysis.

### Gene expression data processing

Gene expression data was downloaded from Interim Analysis 12 (IA12) data release of the Multiple Myeloma Research Foundation. We analyzed the influence of the most highly expressed genes on all other gene values and concluded that special consideration was required to avoid batch effects resulting from highly expressed genes artificially lowering the TPM normalized expression values. In particular, the top 10 most highly expressed transcripts account for more of the mapped reads than the remaining 59,000+ transcripts combined, and the percent of mapped reads attributed to the top 10 genes is strongly anticorrelated to the number of unique transcripts detected (**Fig. S1**). We implemented a custom normalization pipeline in Python that is similar to trimmed mean of M values (TMM) plus quantile normalization (see *miner*.*preprocess*: https://github.com/MattWallScientist/miner3).

### Mutation data processing

Binary mutation matrices were generated such that columns were indexed by patient identifiers, rows were indexed by mutations, and the value of entry (i, j) = 1 if gene *i* was mutated in patient sample *j* and (i, j) = 0 otherwise. The mutation calls used to populate this matrix were taken from the MMRF IA12 clinical data tables provided by the Multiple Myeloma Research Foundation.

### Clinical data processing

Clinical outcomes data was downloaded from the MMRF Researcher Gateway. We used the GuanRank^30^ of time to progression-free survival and normalized the values to fall between 0 and 1. This processing optimizes the value of censored data for regression and classification problems. The code for this normalization is made available at our GitHub page (https://github.com/MattWallScientist/miner3).

### Test dataset acquisition and processing

HOVON65 (GSE19784) and UAMS (GSE24080) datasets were downloaded from NCBI/GEO and processed with the oligo R package to provide RMA normalization. For gene level files with multiple probes mapping to a single gene, log_2_ intensities were combined via the geometric mean. No quantile normalization or mean variance scaling has been computed between studies. The gene expression data as provided was Z-scored and the normalized GuanRank was applied to the progression-free survival data.

### Network inference by Mechanistic Inference of Node-Edge Relationships (MINER)

MINER comprises many functions for the quality control processing, analysis, and predictive model generation from gene expression data in the context of an inferred transcriptional regulatory network (TRN). The TRN is generated by a multi-step pipeline that starts with unsupervised clustering of gene expression, then integrates prior knowledge databases (e.g., transcription factor binding site database), and performs causal inference when the appropriate data (e.g., somatic mutations, copy number variation, etc.) is available. The resulting MINER TRN is composed of units, called regulons, that comprise a set of co-expressed genes sharing a binding site for a regulator whose expression correlates to the first principal component (i.e., the eigengene) of the genes. Additional information, such as upstream causal influences or risk of disease progression as a function of expression level, are associated to each regulon in the network to enable a modular structure. Tutorials of the MINER pipeline and all associated code is available on our GitHub page. The CM TRN presented herein was inferred using the MINER pipeline with the following parameter values: minimum number of genes in a co-expressed set of genes = 6, minimum number of genes in a regulon = 5, minimum magnitude of correlation between regulator and regulon eigengene = 0.2, maximum p-value of binding site enrichment within co-expressed genes = 0.05.

### Calculation of discrete regulon activity

For each normalized sample, the genes were ranked from lowest to highest expression and partitioned into three equal parts: a lower, middle, and top third. Given satisfactory normalization of the gene expression, for example by TMM normalization or the related method proposed in this work, we can form a null hypothesis that a random selection of genes with no co-expressed relationship will tend to distribute evenly between the top, middle, and bottom third of the ranked genes. We can then use a binomial distribution with p = 1/3 to model probability that *k* genes fall into the same third given a selection of *N* genes, where *N* ≥*k*. A default *p-*value of 0.05 is used as a cutoff for rejecting the null hypothesis that the chosen set of genes are not co-expressed. Genes that pass this coherent cutoff in the lower third are labeled “under-expressed”, and those that pass in the upper third all labeled “over-expressed”. All other cases are assigned a label of “neither”. Accordingly, we generate a matrix with values {-1, 0, 1} for the discrete activity of all regulons in all samples.

### Inference of miRNA regulation

The effects of miRNA regulation were inferred by the Framework for Inference of Regulation by MiRNAs (FIRM).^72^ The MINER pipeline can be directly applied to miRNA regulators in the same fashion as for transcription factors, but the lack of miRNA-seq data severely limited the reliable quantitative detection of miRNA transcripts. In this case, MINER defaults to testing for enrichment of miRNA targets in co-expressed clusters but does not enforce correlation of the miRNA expression. FIRM enables enrichment analysis to infer miRNA regulation when co-expressed gene sets are available, but reliable miRNA-seq data is not. We used the default parameters and significance thresholds of *p* = 0.05.

### Risk prediction

We performed Ridge regression (scikit-learn) against the normalized GuanRank of PFS. The MMRF regulon activity was subset to include only the 20% of patients with highest-risk and 50% of patients with lowest risk. The 20% of patients with highest-risk in each dataset were labeled as high-risk and all others were labeled as low-risk for calculation of AUCs. The regularization parameter was selected by randomly splitting the regulon activity into a training and test set, then training a Ridge model on the training set and calculating the AUC of the test set prediction. This was repeated 500 times and the optimized regularization parameter was selected as that which maximized the mean AUC of the 500 tests.

### Validation of univariate risk prediction

The top 100 genes that stratified risk in MMRF IA12 via gene expression were intersected with the top 100 genes that stratified risk via network activity, yielding 39 genes. These 39 genes were evaluated by area under the receiver operating characteristic curve (AUC) using their gene expression or network activity values as predictors of risk in GSE24080 and GSE19784. Random permutations of network activity values were used as a reference for random prediction.

### Association of additional information to causal flows

For each causal flow, we integrated additional information by using a custom pipeline (miner_output_merge.py) that included the following processing steps. *i)* For a given regulon in each causal flow, gene members were collected and queried against OpenTargets database (https://www.targetvalidation.org/) to collect all drugs associated with a given gene for multiple myeloma (get_opentargets.py). *ii*) Similarly, functional enrichment with GO biological process terms (BH-corrected p-value ≤ 0.05) was performed for each regulon (GO_enrichment.R) followed by *iii)* association with Hallmarks of Cancer by using semantic similarity (Lin Semantic Similarity score > 0.4).^17,73,74^ (goSimHallmarksOfCancer.R) *iv)* Putative miRNA regulators via the FIRM pipeline were also associated with each causal flow as described before.

## Acknowledgements

We thank the members of the Baliga lab for critical discussions. Funding was provided by the National Science Foundation (NSF ABI 1565166), National Institute of Health (NIH NIAID R01 AI141953), Institute for Systems Biology, and Celgene/BMS.

## Author Contributions

M.A.W. developed MINER, constructed the MM TRN, analyzed data, interpreted results, prepared the figures, and wrote the manuscript.

A.L.G.L, and S.T. supported the development of MINER, performed the drug mapping analysis, interpreted results, implemented the web portal, prepared the figures, and wrote the manuscript.

W.W. supported the development of MINER, performed the drug mapping analysis, and implemented the web portal.

M.J.M processed the GSE24080 and GSE19784 validation datasets. S.A.D., D.J.R., provided ideas for modeling.

A.P.D. and M.W.B.T. provided guidance on research design and data interpretation.

D.B. provided input into overarching research goals.

R.M.H. conceived the project, formulated overarching research goals.

A.V.R. conceived the project, formulated overarching research goals and aims, designed the research, guided the analysis, and oversaw the project.

N.S.B. conceived the project, formulated overarching research goals and aims, designed the research, guided the analysis, and oversaw the project, interpreted results, prepared figures, and wrote manuscript.

## Conflict-of-interest disclosure

S.A.D., D.L.R., A.P.D., M.W.B.T., D.B. and A.V.R. declare employment and equity ownership for Bristol-Myers Squibb Corporation. R.M.H. declares former employment and equity ownership for Celgene Corporation; membership on an entity’s Board of Directors or advisory committee for Adaptive Biotechnologies and NanoString Technologies; consultancy at Fraizer Healthcare Partners.

## Notes

https://github.com/MattWallScientist/miner3

## References

1. Becker N. Epidemiology of Multiple Myeloma. Multiple Myeloma. 2011;

2. Palumbo A, Anderson K. Multiple Myeloma. N Engl J Med. 2011;364(11):1046–1060.

3. Kumar SK, Rajkumar V, Kyle RA, et al. Multiple myeloma. Nat Rev Dis Primers. 2017;3(1):17046.

4. Manier S, Salem KZ, Park J, et al. Genomic complexity of multiple myeloma and its clinical implications. Nat Rev Clin Oncol. 2017;14(2):100–113.

5. Zhan F, Huang Y, Colla S, et al. The molecular classification of multiple myeloma. Blood. 2006;108(6):2020–2028.

6. Lohr JG, Stojanov P, Carter SL, et al. Widespread Genetic Heterogeneity in Multiple Myeloma: Implications for Targeted Therapy. Cancer Cell. 2014;25(1):91–101.

7. Turesson I, Bjorkholm M, Blimark CH, et al. Rapidly changing myeloma epidemiology in the general population: Increased incidence, older patients, and longer survival. Eur J Haematol. 2018;101(2):237–244.

8. Abramson HN. The Multiple Myeloma Drug Pipeline—2018: A Review of Small Molecules and Their Therapeutic Targets. Clinical Lymphoma Myeloma and Leukemia. 2018;18(9):611–627.

9. Kumar SK, Callander NS, Alsina M, et al. NCCN Guidelines Insights: Multiple Myeloma, Version 3.2018. J Natl Compr Canc Netw. 2018;16(1):11–20.

10. Dimopoulos MA, Richardson PG, Moreau P, Anderson KC. Current treatment landscape for relapsed and/or refractory multiple myeloma. Nature Reviews Clinical Oncology. 2015;12(1):42–54.

11. Kumar SK, Dispenzieri A, Lacy MQ, et al. Continued improvement in survival in multiple myeloma: changes in early mortality and outcomes in older patients. Leukemia. 2014;28(5):1122–1128.

12. Rajkumar SV. Myeloma today: Disease definitions and treatment advances: Myeloma Today. Am. J. Hematol. 2016;91(1):90–100.

13. Robak P, Drozdz I, Szemraj J, Robak T. Drug resistance in multiple myeloma. Cancer Treatment Reviews. 2018;70:199–208.

14. Nass J, Efferth T. Drug targets and resistance mechanisms in multiple myeloma. CDR. 2018;

15. Di Marzo L, Desantis V, Solimando AG, et al. Microenvironment drug resistance in multiple myeloma: emerging new players. Oncotarget. 2016;7(37):.

16. Nijhof IS, van de Donk NWCJ, Zweegman S, Lokhorst HM. Current and New Therapeutic Strategies for Relapsed and Refractory Multiple Myeloma: An Update. Drugs. 2018;78(1):19–37.

17. Plaisier CL, O’Brien S, Bernard B, et al. Causal Mechanistic Regulatory Network for Glioblastoma Deciphered Using Systems Genetics Network Analysis. Cell Systems. 2016;3(2):172–186.

18. Reiss DJ, Baliga NS, Bonneau R. Integrated biclustering of heterogeneous genome-wide datasets for the inference of global regulatory networks. BMC Bioinformatics. 2006;7(1):280.

19. Bonneau R, Reiss DJ, Shannon P, et al. The Inferelator: an algorithm for learning parsimonious regulatory networks from systems-biology data sets de novo. Genome Biol. 2006;7(5):R36.

20. Brooks AN, Reiss DJ, Allard A, et al. A system-level model for the microbial regulatory genome. Mol Syst Biol. 2014;10(7):740.

21. Margolin AA, Nemenman I, Basso K, et al. ARACNE: An Algorithm for the Reconstruction of Gene Regulatory Networks in a Mammalian Cellular Context. BMC Bioinformatics. 2006;7(S1):S7.

22. Agnelli L, Forcato M, Ferrari F, et al. The Reconstruction of Transcriptional Networks Reveals Critical Genes with Implications for Clinical Outcome of Multiple Myeloma. Clinical Cancer Research. 2011;17(23):7402–7412.

23. Laganà A, Perumal D, Melnekoff D, et al. Integrative network analysis identifies novel drivers of pathogenesis and progression in newly diagnosed multiple myeloma. Leukemia. 2018;32(1):120–130.

24. Liu Y, Yu H, Yoo S, et al. A Network Analysis of Multiple Myeloma Related Gene Signatures. Cancers. 2019;11(10):1452.

25. Robinson MD, Oshlack A. A scaling normalization method for differential expression analysis of RNA-seq data. Genome Biol. 2010;11(3):R25.

26. Shi L, Campbell G, Jones WD, et al. The MicroArray Quality Control (MAQC)-II study of common practices for the development and validation of microarray-based predictive models. Nature Biotechnology. 2010;28(8):827–838.

27. Broyl A, Hose D, Lokhorst H, et al. Gene expression profiling for molecular classification of multiple myeloma in newly diagnosed patients. Blood. 2010;116(14):2543–2553.

28. Strimmer K. A unified approach to false discovery rate estimation. BMC Bioinformatics. 2008;9(1):303.

29. Klaus B, Reisenauer S. An end to end workflow for differential gene expression using Affymetrix microarrays [version 2; peer review: 2 approved]. F1000Research. 2018;5(1384):.

30. Huang Z, Zhang H, Boss J, et al. Complete hazard ranking to analyze right-censored data: An ALS survival study. PLoS Comput Biol. 2017;13(12):e1005887.

31. Neri P, Ren L, Gratton K, et al. Bortezomib-induced “BRCAness” sensitizes multiple myeloma cells to PARP inhibitors. Blood. 2011;118(24):6368–6379.

32. Gu C, Yang Y, Sompallae R, et al. FOXM1 is a therapeutic target for high-risk multiple myeloma. Leukemia. 2016;30(4):873–882.

33. Shaughnessy JD, Zhan F, Burington BE, et al. A validated gene expression model of highrisk multiple myeloma is defined by deregulated expression of genes mapping to chromosome 1. Blood. 2007;109(6):2276–2284.

34. Kuiper R, Broyl A, de Knegt Y, et al. A gene expression signature for high-risk multiple myeloma. Leukemia. 2012;26(11):2406–2413.

35. Hose D, Reme T, Hielscher T, et al. Proliferation is a central independent prognostic factor and target for personalized and risk-adapted treatment in multiple myeloma. Haematologica. 2011;96(1):87–95.

36. Mason MJ, Schinke C, Eng CLP, et al. Multiple Myeloma DREAM Challenge Reveals Epigenetic Regulator *PHF19* As Marker of Aggressive Disease. Cancer Biology; 2019.

37. Resnitzky D, Reed SI. Different roles for cyclins D1 and E in regulation of the G1-to-S transition. Mol. Cell. Biol. 1995;15(7):3463–3469.

38. Pawlyn C, Morgan GJ. Evolutionary biology of high-risk multiple myeloma. Nature Reviews Cancer. 2017;17(9):543–556.

39. Anders L, Ke N, Hydbring P, et al. A Systematic Screen for CDK4/6 Substrates Links FOXM1 Phosphorylation to Senescence Suppression in Cancer Cells. Cancer Cell. 2011;20(5):620–634.

40. Xia C, Bao Z, Tabassam F, et al. GCIP, a Novel Human Grap2 and Cyclin D Interacting Protein, Regulates E2F-mediated Transcriptional Activity. J. Biol. Chem. 2000;275(27):20942–20948.

41. Ma W, Stafford LJ, Li D, et al. GCIP/CCNDBP1, a helix–loop–helix protein, suppresses tumorigenesis. Journal of Cellular Biochemistry. 2007;100(6):1376–1386.

42. Kronke J, Udeshi ND, Narla A, et al. Lenalidomide Causes Selective Degradation of IKZF1 and IKZF3 in Multiple Myeloma Cells. Science. 2014;343(6168):301–305.

43. Dong Y, Poellinger L, Gustafsson J-Å, Okret S. Regulation of Glucocorticoid Receptor Expression: Evidence for Transcriptional and Posttranslational Mechanisms. Molecular Endocrinology. 1988;2(12):1256–1264.

44. Oerlemans R, Franke NE, Assaraf YG, et al. Molecular basis of bortezomib resistance: proteasome subunit β5 (PSMB5) gene mutation and overexpression of PSMB5 protein. Blood. 2008;112(6):2489–2499.

45. Reale A, Khong T, Mithraprabhu S, et al. TOP2A knockdown resensitizes carfilzomib-resistant HMCLs to carfilzomib. Clinical Lymphoma, Myeloma and Leukemia. 2015;15:e68.

46. Franqui-Machin R, Hao M, Bai H, et al. Destabilizing NEK2 overcomes resistance to proteasome inhibition in multiple myeloma. Journal of Clinical Investigation. 2018;128(7):2877–2893.

47. Turner JG, Dawson JL, Grant S, et al. Treatment of acquired drug resistance in multiple myeloma by combination therapy with XPO1 and topoisomerase II inhibitors. J Hematol Oncol. 2016;9(1):73.

48. Punnoose EA, Leverson JD, Peale F, et al. Expression Profile of BCL-2, BCL-XL, and MCL-1 Predicts Pharmacological Response to the BCL-2 Selective Antagonist Venetoclax in Multiple Myeloma Models. Molecular Cancer Therapeutics. 2016;15(5):1132–1144.

49. Kumar S, Kaufman JL, Gasparetto C, et al. Efficacy of venetoclax as targeted therapy for relapsed/refractory t(11;14) multiple myeloma. Blood. 2017;130(22):2401–2409.

50. Richardson PG, Bringhen S, Voorhees P, et al. Melflufen plus dexamethasone in relapsed and refractory multiple myeloma (O-12-M1): a multicentre, international, open-label, phase 1–2 study. The Lancet Haematology. 2020;

51. Botta C, GullÃ A, Correale P, Tagliaferri P, Tassone P. Myeloid-Derived Suppressor Cells in Multiple Myeloma: Pre-Clinical Research and Translational Opportunities. Front. Oncol. 2014;4:.

52. Mohan M, Weinhold N, Schinke C, et al. Daratumumab in high-risk relapsed/refractory multiple myeloma patients: adverse effect of chromosome 1q21 gain/amplification and GEP70 status on outcome. Br J Haematol. 2020;189(1):67–71.

53. Kelley N, Jeltema D, Duan Y, He Y. The NLRP3 Inflammasome: An Overview of Mechanisms of Activation and Regulation. IJMS. 2019;20(13):3328.

54. Nakamura K, Kassem S, Cleynen A, et al. Dysregulated IL-18 Is a Key Driver of Immunosuppression and a Possible Therapeutic Target in the Multiple Myeloma Microenvironment. Cancer Cell. 2018;33(4):634–648.e5.

55. Roy P, Sarkar U, Basak S. The NF-κB Activating Pathways in Multiple Myeloma. Biomedicines. 2018;6(2):59.

56. Wang S, Song R, Wang Z, et al. S100A8/A9 in Inflammation. Front. Immunol. 2018;9:1298.

57. Bianchi G, Munshi NC. Pathogenesis beyond the cancer clone(s) in multiple myeloma. Blood. 2015;125(20):3049–3058.

58. Terpos E, Ntanasis-Stathopoulos I, Gavriatopoulou M, Dimopoulos MA. Pathogenesis of bone disease in multiple myeloma: from bench to bedside. Blood Cancer Journal. 2018;8(1):7.

59. Hideshima T, Chauhan D, Schlossman R, Richardson P, Anderson KC. The role of tumor necrosis factor a in the pathophysiology of human multiple myeloma: therapeutic applications. 9.

60. Oldenborg P-A, Zheleznyak A, Fang Y-F, et al. Role of CD47 as a Marker of Self on Red Blood Cells. Science. 2000;288(5473):2051.

61. Jaiswal S, Jamieson CHM, Pang WW, et al. CD47 Is Upregulated on Circulating Hematopoietic Stem Cells and Leukemia Cells to Avoid Phagocytosis. Cell. 2009;138(2):271–285.

62. Hobo W, Norde WJ, Schaap N, et al. B and T Lymphocyte Attenuator Mediates Inhibition of Tumor-Reactive CD8 ^+^ T Cells in Patients After Allogeneic Stem Cell Transplantation. J.I. 2012;189(1):39–49.

63. Huang GN, Huso DL, Bouyain S, et al. NFAT Binding and Regulation of T Cell Activation by the Cytoplasmic Scaffolding Homer Proteins. Science. 2008;319(5862):476–481.

64. Seckinger A, Meiβner T, Moreaux J, et al. Clinical and prognostic role of annexin A2 in multiple myeloma. Blood. 2012;120(5):1087–1094.

65. Zingoni A, Cecere F, Vulpis E, et al. Genotoxic Stress Induces Senescence-Associated ADAM10-Dependent Release of NKG2D MIC Ligands in Multiple Myeloma Cells. J.I. 2015;195(2):736–748.

66. Xiao G, Wang X, Sheng J, et al. Soluble NKG2D ligand promotes MDSC expansion and skews macrophage to the alternatively activated phenotype. J Hematol Oncol. 2015;8(1):13.

67. Doubrovina ES, Doubrovin MM, Vider E, et al. Evasion from NK Cell Immunity by MHC Class I Chain-Related Molecules Expressing Colon Adenocarcinoma. J Immunol. 2003;171(12):6891–6899.

68. Groh V, Wu J, Yee C, Spies T. Tumour-derived soluble MIC ligands impair expression of NKG2D and T-cell activation. Nature. 2002;419(6908):734–738.

69. Märten A, von Lilienfeld-Toal M, Büchler MW, Schmidt J. Soluble MIC is elevated in the serum of patients with pancreatic carcinoma diminishing γδ T cell cytotoxicity. Int. J. Cancer. 2006;119(10):2359–2365.

70. Zahoor M, Westhrin M, Aass KR, et al. Hypoxia promotes IL-32 expression in myeloma cells, and high expression is associated with poor survival and bone loss. Blood Advances. 2017;1(27):2656–2666.

71. Lin X, Yang L, Wang G, et al. Interleukin-32α promotes the proliferation of multiple myeloma cells by inducing production of IL-6 in bone marrow stromal cells. Oncotarget. 2017;8(54):.

72. Plaisier CL, Pan M, Baliga NS. A miRNA-regulatory network explains how dysregulated miRNAs perturb oncogenic processes across diverse cancers. Genome Research. 2012;22(11):2302–2314.

73. Hanahan D, Weinberg RA. Hallmarks of Cancer: The Next Generation. Cell. 2011;144(5):646–674.

74. Knijnenburg TA, Bismeijer T, Wessels LFA, Shmulevich I. A multilevel pan-cancer map links gene mutations to cancer hallmarks. Chin J Cancer. 2015;34(3):48.

