## Supplementary Information for "Genetic program activity delineates risk, relapse, and therapy responsiveness in Multiple Myeloma"

| Gene | Other | Del 17 | Amp 1q | t(11;14) | t(4;14) | t(14;16) | MYC |
|---|---|---|---|---|---|---|---|
| CRBN | -0.12 | -- | -0.14 | -- | -0.16 | -- | -- |
| IKZF1 | -- | -- | -- | -- | -0.21 | -- | -- |
| IKZF3 | -- | -- | -- | -- | -- | -0.38 | -- |
| NR3C1 | -- | -- | -0.12 | -- | -- | -0.40 | -- |
| PSMB5 | -0.10 | -- | -- | -- | 0.19 | -- | -- |
| HDAC1 | -- | -- | -- | -- | 0.18 | -- | -- |
| HDAC2 | 0.10 | -- | 0.14 | -- | 0.32 | 0.30 | -- |
| HDAC6 | -- | -- | -0.16 | -- | -0.18 | -- | -- |
| HDAC9 | -0.10 | -- | -- | -- | -- | -- | -- |
| APH1A | 0.17 | 0.55 | 0.16 | -- | 0.39 | -- | -- |
| PARP1 | 0.14 | 0.36 | 0.15 | -- | 0.44 | -- | -- |
| KIF11 | 0.16 | 0.32 | 0.32 | 0.24 | 0.37 | -- | -- |
| GSK3B | 0.12 | 0.32 | 0.15 | -- | 0.25 | -- | -- |
| AURKA | 0.13 | -- | 0.24 | 0.17 | 0.43 | -- | -- |
| AURKB | 0.15 | 0.31 | 0.28 | 0.21 | 0.41 | -- | -- |
| ANPEP | 0.13 | -- | -- | -- | -- | 0.49 | -- |

**Table S1. Significant correlations between risk of disease progression and network activity for targets of MM therapies.** The spearman correlation coefficient is listed for all results satisfying $p < 0.1$.

| Gene | Other | Del 17 | Amp 1q | t(11;14) | t(4;14) | t(14;16) | MYC |
|---|---|---|---|---|---|---|---|
| CRBN | -0.10 | -- | -0.13 | -- | -0.31 | -- | -- |
| IKZF1 | -- | -- | -- | -- | -- | -- | -- |
| IKZF3 | -- | -- | -- | -- | -- | -- | -- |
| NR3C1 | -0.11 | -- | -0.12 | -- | -0.17 | -0.56 | -- |
| PSMB5 | -- | -- | -- | -- | -- | -- | -- |
| HDAC1 | -- | -- | -- | -- | 0.17 | -- | -- |
| HDAC2 | -- | -- | 0.12 | -- | 0.27 | 0.31 | -- |
| HDAC6 | -- | -- | -- | -- | -- | -- | -- |
| HDAC9 | -- | -- | -0.17 | -- | -0.25 | -- | -0.28 |
| APH1A | 0.09 | 0.31 | -- | 0.17 | 0.24 | -- | 0.29 |
| PARP1 | 0.00 | -- | 0.13 | -- | 0.33 | -- | -- |
| KIF11 | 0.13 | 0.35 | 0.25 | 0.24 | 0.38 | 0.32 | 0.33 |
| GSK3B | -- | -- | -- | -- | -- | -- | -- |
| AURKA | -- | 0.37 | 0.22 | -- | 0.40 | -- | -- |
| AURKB | 0.12 | 0.34 | 0.28 | 0.27 | 0.36 | -- | -- |
| ANPEP | 0.14 | -- | -- | -- | -- | -- | -- |

**Table S2. Significant correlations between risk of disease progression and gene expression for targets of MM therapies.** The spearman correlation coefficient is listed for all results satisfying $p < 0.1$.

| Gene | Other | Del 17 | Amp 1q | t(11;14) | t(4;14) | t(14;16) | MYC |
|---|---|---|---|---|---|---|---|
| CRBN | 3.02E-02 | 3.92E-01 | 4.55E-02 | 2.09E-01 | 1.00E-01 | 7.59E-01 | 5.69E-01 |
| IKZF1 | 4.36E-01 | 8.36E-01 | 2.42E-01 | 3.51E-01 | 3.48E-02 | 2.16E-01 | 8.54E-01 |
| IKZF3 | 1.10E-01 | 7.11E-01 | 1.08E-01 | 8.88E-01 | 6.89E-01 | 2.08E-02 | 3.52E-01 |
| NR3C1 | 1.60E-01 | 7.62E-01 | 9.50E-02 | 7.72E-01 | 7.87E-01 | 1.46E-02 | 5.79E-01 |
| PSMB5 | 7.09E-02 | 8.95E-01 | 9.63E-01 | 6.46E-01 | 4.88E-02 | 4.11E-01 | 8.46E-01 |
| HDAC1 | 7.72E-01 | 3.26E-01 | 3.24E-01 | 5.26E-01 | 7.21E-02 | 4.30E-01 | 7.92E-01 |
| HDAC2 | 7.63E-02 | 1.69E-01 | 4.54E-02 | 2.47E-01 | 7.64E-04 | 6.74E-02 | 3.14E-01 |
| HDAC6 | 9.50E-01 | 8.01E-01 | 2.39E-02 | 6.35E-01 | 6.21E-02 | 3.04E-01 | 5.60E-01 |
| HDAC9 | 7.18E-02 | 3.90E-01 | 1.07E-01 | 4.33E-01 | 2.64E-01 | 5.30E-01 | 4.91E-01 |
| APH1A | 1.92E-03 | 6.40E-04 | 1.92E-02 | 8.53E-01 | 4.76E-05 | 5.73E-01 | 2.98E-01 |
| PARP1 | 1.16E-02 | 3.36E-02 | 3.75E-02 | 3.22E-01 | 3.49E-06 | 8.45E-01 | 7.11E-02 |
| KIF11 | 4.53E-03 | 6.36E-02 | 4.30E-06 | 2.69E-03 | 9.96E-05 | 2.14E-01 | 1.29E-01 |
| GSK3B | 2.70E-02 | 6.14E-02 | 3.14E-02 | 3.62E-01 | 8.77E-03 | 8.58E-01 | 2.50E-01 |
| AURKA | 1.81E-02 | 1.27E-01 | 4.58E-04 | 3.66E-02 | 5.61E-06 | 1.48E-01 | 2.02E-01 |
| AURKB | 5.56E-03 | 7.02E-02 | 3.85E-05 | 9.34E-03 | 1.57E-05 | 1.39E-01 | 2.04E-01 |
| ANPEP | 1.73E-02 | 9.03E-01 | 4.91E-01 | 5.28E-01 | 9.96E-01 | 2.30E-03 | 4.53E-01 |

**Table S3. Spearman correlation *p*-values of risk of disease progression versus network activity for targets of MM therapies.** Risk of disease progression was quantified by the GuanRank algorithm.

| Gene | Other | Del 17 | Amp 1q | t(11;14) | t(4;14) | t(14;16) | MYC |
|---|---|---|---|---|---|---|---|
| CRBN | 7.81E-02 | 4.49E-01 | 6.01E-02 | 1.28E-01 | 1.49E-03 | 3.86E-01 | 9.84E-01 |
| IKZF1 | 2.45E-01 | 1.11E-01 | 8.77E-01 | 6.24E-01 | 8.66E-01 | 4.89E-01 | 5.58E-01 |
| IKZF3 | 2.68E-01 | 3.50E-01 | 2.63E-01 | 8.57E-01 | 6.13E-01 | 6.89E-01 | 1.57E-01 |
| NR3C1 | 4.80E-02 | 6.19E-01 | 8.37E-02 | 6.18E-01 | 7.86E-02 | 3.37E-04 | 1.67E-01 |
| PSMB5 | 6.85E-01 | 3.09E-01 | 9.41E-01 | 8.98E-01 | 4.12E-01 | 8.76E-01 | 6.70E-01 |
| HDAC1 | 4.08E-01 | 7.88E-01 | 3.66E-01 | 9.99E-01 | 7.47E-02 | 1.41E-01 | 6.51E-01 |
| HDAC2 | 8.98E-01 | 3.12E-01 | 8.04E-02 | 3.69E-01 | 5.59E-03 | 6.20E-02 | 5.80E-01 |
| HDAC6 | 8.15E-01 | 4.63E-01 | 9.48E-01 | 3.39E-01 | 6.60E-01 | 6.57E-01 | 2.51E-01 |
| HDAC9 | 4.61E-01 | 4.40E-01 | 1.29E-02 | 6.84E-01 | 1.01E-02 | 1.80E-01 | 9.72E-02 |
| APH1A | 8.85E-02 | 6.99E-02 | 1.91E-01 | 3.89E-02 | 1.22E-02 | 2.78E-01 | 8.43E-02 |
| PARP1 | 3.89E-01 | 4.72E-01 | 6.35E-02 | 7.90E-01 | 6.77E-04 | 6.17E-01 | 3.51E-01 |
| KIF11 | 2.25E-02 | 3.72E-02 | 2.59E-04 | 2.76E-03 | 7.14E-05 | 5.53E-02 | 4.84E-02 |
| GSK3B | 2.42E-01 | 6.44E-01 | 7.14E-01 | 4.27E-01 | 2.11E-01 | 1.12E-01 | 6.48E-01 |
| AURKA | 2.07E-01 | 2.82E-02 | 1.69E-03 | 1.02E-01 | 2.13E-05 | 1.36E-01 | 1.13E-01 |
| AURKB | 3.35E-02 | 4.89E-02 | 4.23E-05 | 8.50E-04 | 1.54E-04 | 2.12E-01 | 1.22E-01 |
| ANPEP | 1.35E-02 | 1.76E-01 | 1.73E-01 | 6.45E-01 | 4.43E-01 | 5.18E-01 | 1.96E-01 |

**Table S4. Spearman correlation *p*-values of risk of disease progression versus gene expression for targets of MM therapies.** Risk of disease progression was quantified by the GuanRank algorithm.

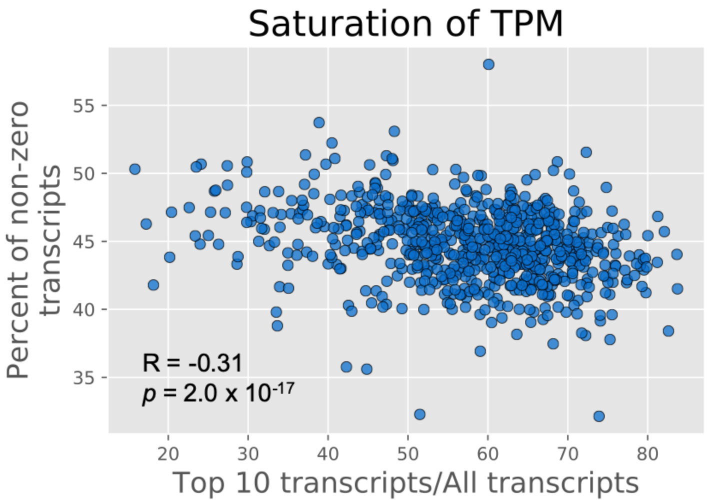


**Figure S1. Unwanted influence of highly-expressed genes on expression data.** The percentage of transcripts that were detected (i.e., non-zero counts) via RNA-seq strongly anti-correlated with the percent of counts attributed to the 10 most highly expressed genes in any sample. This suggests that the highly-expressed genes are introducing bias into the experiments by saturating the reads and thus under-representing the expression of all other transcripts. This effect must be accounted for during pre-processing of the RNA-seq data by an approach such as TMM normalization that is not biased by outlier expression.

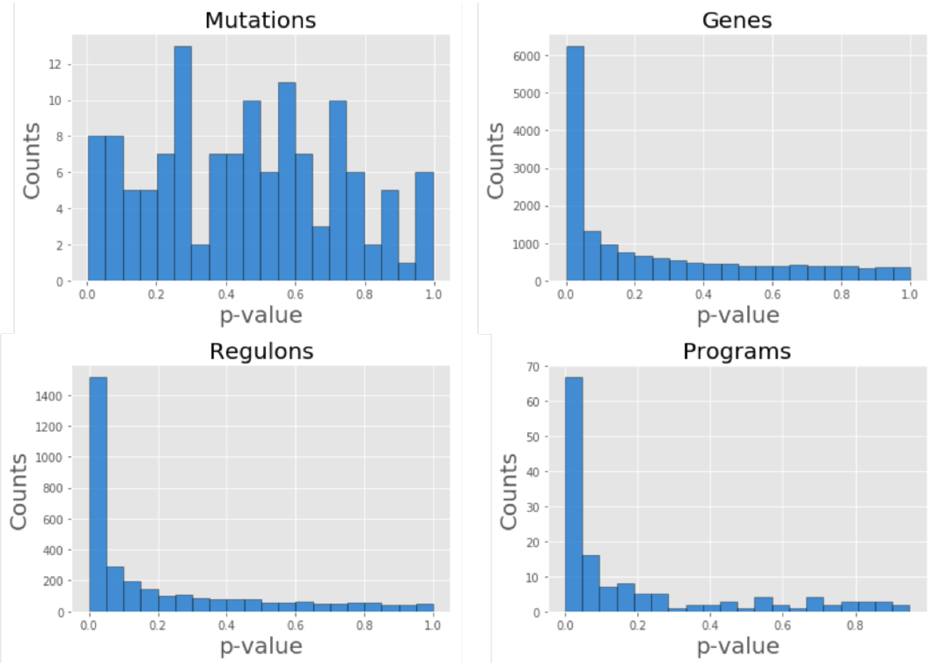


**Figure S2. Distribution of uncorrected p-values for various predictive features.** The distribution of uncorrected p-values gives an indication of predictive power via setting expectations for false discovery rate. If the proportion of features with $p<0.1$ is significantly greater than 0.1, then the feature tends to stratify better than by random chance. Here we see that mutations do not stratify better than random chance, but that programs, regulons, and gene expression do.

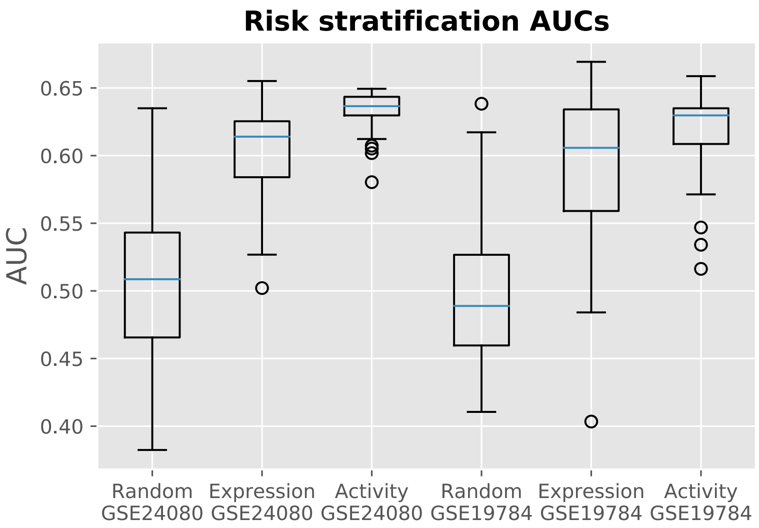


**Figure S3. Network activity improves univariate risk prediction.** The individual genes most predictive of risk in MMRF IA12 were evaluated for their ability to stratify risk in the validation datasets (GSE24080 and GSE19784). In both cases the network activity outperformed the gene expression.

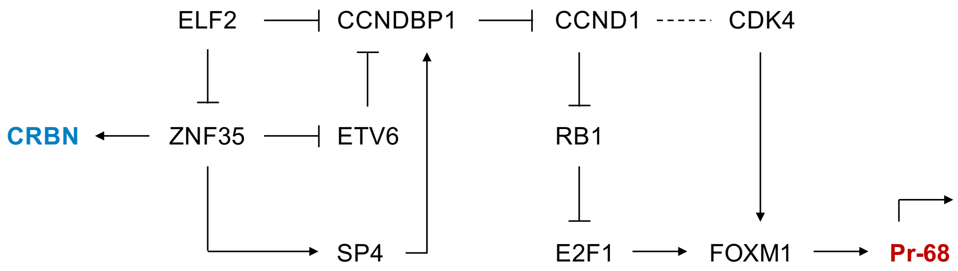


**Figure S4. Regulatory circuit connects CRBN and CCNDBP1 in CM TRN of MM.** The activity of CRBN and CCNDBP1 are inferred to correlate due to shared upstream regulators of ELF2 and ZNF35. The link between CCNDBP1 and the CCND1-CDK4 complex connects this circuit to Pr-68.

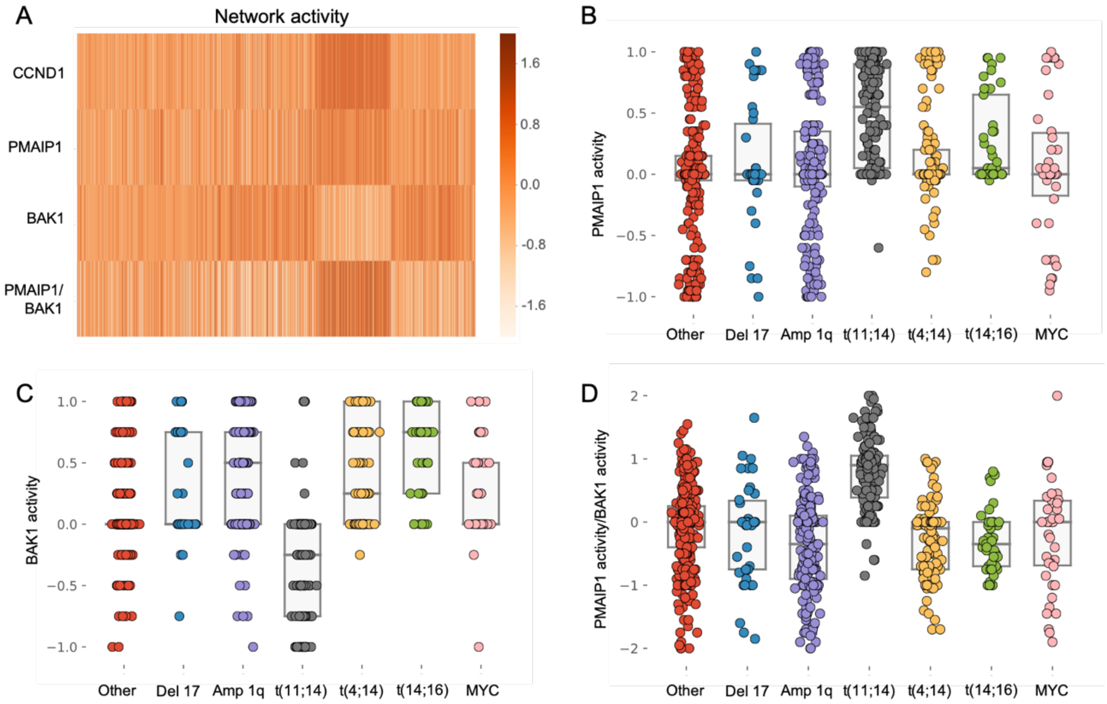


**Figure S5. Subtype-specific network activity profiles of BCL2-family intrinsic apoptosis genes.** The network-constrained activity of genes in the BCL2 family that regulate intrinsic apoptosis signaling exhibit distinct profiles. **A,** The genes PMAIP1 and BAK1 are particularly noteworthy for their changes in activity upon high CCND1 expression (e.g., as occurs in t(11;14) patients). **B-D,** The network activity of PMAIP1 (B) and BAK1 (C) exhibit significantly different activity levels in t(11;14) patients in comparison to other subtypes, and the ratio of PMAIP1/BAK1 (D) is especially high in these patients.
